# An H3-K9-me-independent binding of Swi6/HP1 to siRNA-DNA hybrids initiates heterochromatin assembly at cognate dg-dh repeats in Fission Yeast

**DOI:** 10.1101/2020.10.21.349050

**Authors:** Jyotsna Kumar, Swati Haldar, Neelima Gupta, Viney Kumar, Manisha Thakur, Keerthivasan Raanin Chandradoss, Debarghya Ghose, Dipak Dutta, Kuljeet Singh Sandhu, Jagmohan Singh

## Abstract

Canonically, heterochromatin formation in fission yeast and metazoans is initiated by di/trimethylation of histone H3 at lysine 9 position by the histone methyltransferase Suv39H1/Clr4, followed by binding of Swi6/HP1 to H3-K9-me2/me3 via its chromodomain. Subsequent self-association of Swi6/HP1 on adjacent nucleosomes or a cooperative interaction between Clr4 and Swi6/HP1 leads to folded heterochromatin structure. HP1 binding to RNA is shown to facilitate its localization at and assembly of heterochromatin in metazoans. Likewise, recruitment of Swi6/HP1 to centromere depends on the RNAi pathway in fission yeast; paradoxically, Swi6/HP1 is also thought to play a role in RNA turnover. Here we provide evidence in support of RNAi-independent recruitment of Swi6. We show that, apart from the low affinity binding to RNAs through its hinge domain, as already reported, Swi6/HP1 displays a hierarchy of increasing binding affinity through its chromodomain to the siRNAs corresponding to specific dg-dh repeats and even stronger binding to the cognate siRNA-DNA hybrids than to the siRNA precursors or general RNAs. Our results support a mechanism of recruitment of Swi6, which is dependent on its specific and high affinity binding to siRNA-DNA hybrid at the dg-dh repeats. This binding, which is independent of, albeit augmented by binding to H3-K9-Me2, leads to heterochromatin formation and silencing. We suggest that the net role of Swi6 in RNA physiology may be regulated by a balance between abundance and affinity of Swi6 towards heterochromatic and euchromatic RNAs and siRNAs.

## Introduction

According to the histone-code hypothesis, eukaryotic chromatin is organized into expressed/euchromatic and repressed/heterochromatic regions through a combinatorial set of distinct post-translational histone modifications (Jenuwein & Allis, 2001). In fission yeast (*Schizosaccharomyces pombe*) and metazoans, freshly assembled chromatin contains histone H3 acetylated at K9 and K14 residues (Nakayama *et al*, 2001; Grewal & Jia, 2007). At heterochromatin regions, the H3-acetyl groups are removed by the deacetylases, Clr3, Clr6 and Sir2, followed by di- and tri-methylation of K9 by Suv39/Clr4 (Nakayama *et al*, 2001). Heterochromatin protein Swi6/HP1 binds to Me2/Me3-H3-K9 through its chromodomain (Bannister *et al*, 2001). Subsequent self-association of HP1/Swi6 through its chromo-shadow domain results in a folded, transcriptionally inactive heterochromatin. An alternative model has invoked a mutually cooperative recruitment of Swi6/HP1 and Clr4/Suv3-9 in heterochromatin formation and spreading (Haldar *et al*, 2011; Al-Sady *et al*, 2013).

The RNAi pathway is also involved in heterochromatin assembly as the *rnai* mutants show impaired H3K9 dimethylation and Swi6 recruitment at the mating type and pericentromeric regions (Volpe *et al*, 2002). According to the current model, the double-stranded (ds) RNAs formed by bidirectional transcription from the *dg-dh* repeat regions are cleaved by Dcr1 to produce the ds siRNAs. These siRNAs are bound to the ARC (<u>A</u>rgonaute small interfering <u>R</u>NA <u>c</u>haperone) complex, which comprises the endoribonuclease 1 (Ago1), Arb1 and Arb2 subunits (Holoch and Moazed, 2015). After the passenger strand release, the guide strand is transferred to the chromatin-bound complex, called RNA-induced transcriptional silencing (RITS) complex (Verdel *et al*, 2004), which consists of Ago1, Chp1-a chromodomain protein and Tas3. The binding of Chp1 to Me2/Me3-K9-H3 tethers the RITS complex to heterochromatin. Subsequently, Rdp1 (RNA-dependent RNA polymerase) copies the non-coding ncRNA strand synthesized by RNA PolII, using the guide strand as a primer, to generate the double-stranded (ds) RNA (Sugiyama *et al*, 2005). In turn, the ds RNA, is cleaved by Dcr1 to produce the siRNAs, which start the new amplification cycle (Martienssen and Moazed, 2015). The RNAi link to the formation of Me2/Me3-K9-H3 may possibly occur through Stc1-dependent recruitment of the histone methyltransferase Clr4-containing CLRC (CLR like Clr4 complex; Hong *et al*, 2005; Bayne *et al*, 2010).

As mentioned above, RNAi pathway in *S. pombe* is linked to recruitment of Swi6/HP1 at the pericentromeric regions. These regions contain the *dg*-*dh* repeats, which are also present at the *cenH* region spanning the *mat2* and *mat3* interval and the subtelomeric genes *tlh1* and *tlh2* (Grewal & Klar, 1997; Hansen *et al*, 2006). However, the exact mechanism by which the RNAi pathway is involved in heterochromatin recruitment of Swi6/HP1 is not known.

In this regard, earlier studies showed that apart from binding to Me2/Me3-Lys9-H3, the chromodomain motif, which is present in Swi6/HP1, also shows strong RNA-binding (Akhtar *et al*, 2000). Furthermore, RNA binding by the murine HP1 through its chromo- and hinge domains is required for heterochromatin assembly (Muchardt *et al*, 2002). However, another study claimed that heterochromatin-transcribed RNA competes with Lys9 methylated H3 for binding to Swi6 and is carried to the exosome (Keller *et al*, 2012).

Here, we demonstrate that Swi6/HP1 binds with high affinity to the *dg*-*dh* specific siRNAs engendered by RNAi machinery-mediated processing of the precursor RNAs and even greater affinity to the cognate RNA-DNA hybrids. This binding is significantly stronger than and qualitatively different from that reported for GFP RNA and euchromatic RNAs (Keller et al., 2012). We also suggest that this mode of binding could facilitate a self-autonomous mode of recruitment of Swi6/HP1 to heterochromatin, which is independent of but may be supplemented by binding to Me2/Me3-K9-H3. We suggest a dynamic role of Swi6/HP1 in harnessing its high affinity binding to the *dg-dh* specific siRNAs to facilitate its recruitment to cognate sites in DNA, while the relatively lower affinity binding of Swi6 through its hinge domain to non-heterochromatic RNAs may be instrumental in regulating turnover of excessive levels of normal or abundant RNAs.

## Results

### Specific binding of Swi6/HP1 to the dg-dh- specific siRNAs

We investigated whether, similar to mouse HP1 (Muchardt et al., 2002), Swi6/HP1 may be recruited to the *dg*-*dh* repeats by binding to the cognate si RNAs in *S. pombe*. This scenario is consistent with the loss of heterochromatin recruitment of Swi6 as well as siRNA generation in *rnai* mutants (Volpe *et al*, 2002). Indeed, several siRNAs have been reported to occur *in vivo* (Reinhardt and Bartel, 2002), or generated by Dicer cleavage of precursor RNA precursor *in vitro* (Djupedal *et al,* 2009). Both these populations of siRNAs were found to map to heterochromatin loci- in particular, the *dg* and *dh* repeats that are shared among the outer repeats (*otr*) on all three centromeres (Djupedal *et al,* 2009), the *cenH* (Grewal & Klar, 1997) region (*mat2-mat3* interval) on *chrII* and the subtelomeric *tlh1* and *tlh2* genes on *chrI* and *chrII* (Hansen *et al*, 2006) (Figure 1A and B; Supplementary Figures A1 and S2; Tables S1 and S2).

**Figure 1.**
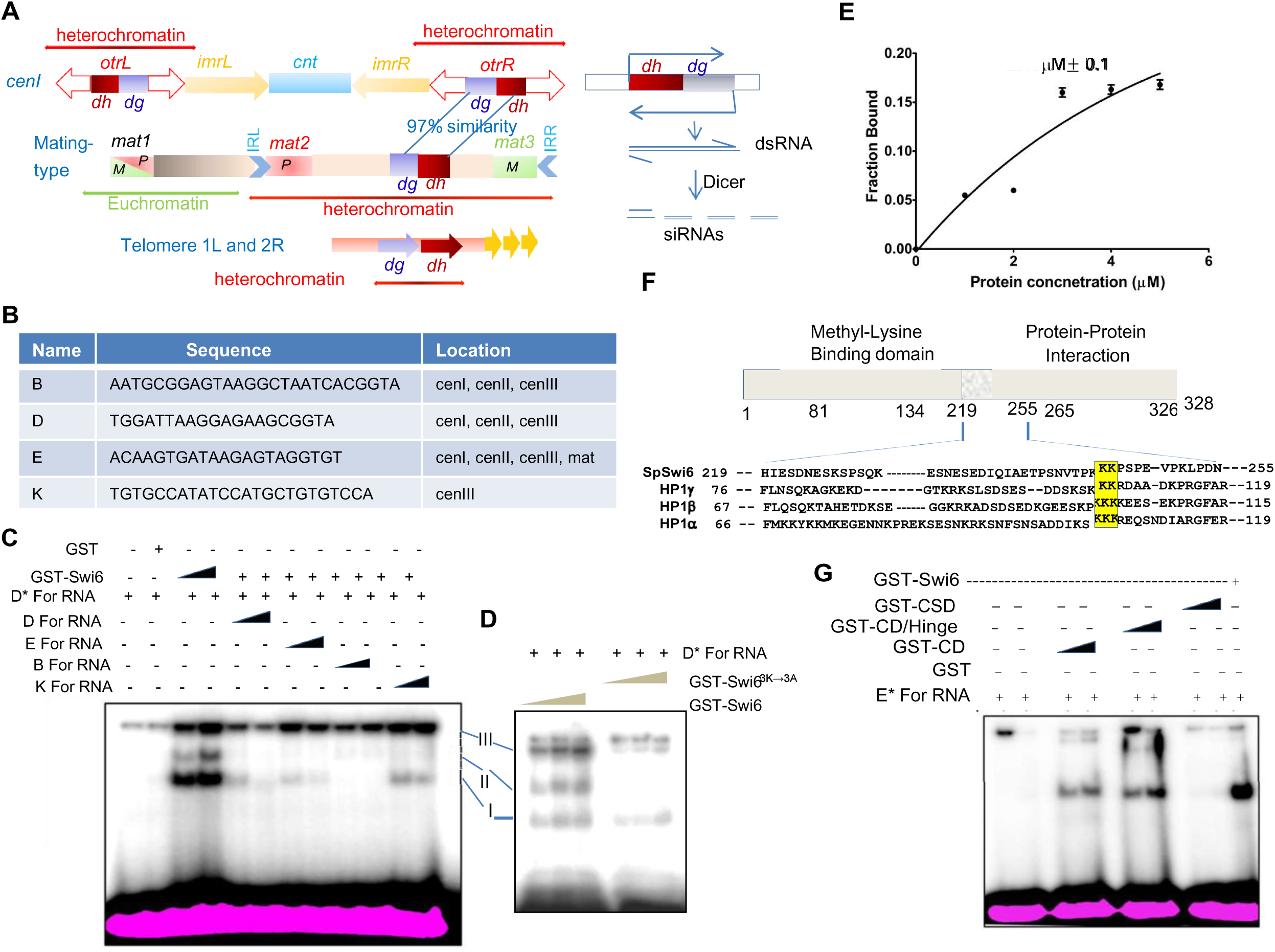
Sequence-specific binding of Swi6 to the *dg*-*dh-*specific siRNAs *in vitro*. A Schematic representation of the *dg* and *dh* regions in the centromere, mating type (*cenH*) and the sub-telomeric genes. *tlh1* and *tlh2*. Right panel, a schematic showing siRNAs being generated from the *dg*-*dh* sequences. B List of si RNA sequences, reported earlier (Reinhardt & Bartel, 2002), that were synthesized *in vitro*. C EMSA assay showing sequence specific binding of Swi6 to the ss D-For RNA and its competition by excess col RNA sequences B, D, E and K. Arrows indicate the complexes I, II and II between Swi6 and ‘D-For’ RNA. D The mutant Swi6^3K→3A^ shows weaker binding to the D-For RNA. EMSA was performed as in (C). E Estimation of equilibrium binding constant (Kd) for the binding of Swi6 to D-For RNA. The data of protein concentration of Swi6 along X axis and the fraction of siRNA bound was plotted as described (Heffler *et al*, 2012). F Schematic depiction of the domain structure of Swi6 showing the chromo-, chromo-hinge and chromo-shadow domains. Alignment of the sequences of the hinge domain between HP1α, HP1β, HP1γ and Swi6, showing conservation of the lysine triplet located at residues 242-244. G EMSA showing that the RNA binding could be ascribed primarily to the CD and CD-hinge but not the CSD.

In accordance with the earlier report showing binding of murine HP1 to RNAs (Muchardt *et al*, 2002), we checked whether Swi6 shows binding to siRNAs. Accordingly, we selected siRNAs found in vivo, labelled A-K (Reinhardt and Bartel, 2002) and produced in vitro (Djupedal *et al*., 2009). By pairwise alignment, we noticed significant sequence conservation among the siRNA sequences (Supplementary Figure 2). We focused on the siRNAs denoted as B, D, E H and K (Reinhardt & Bartel, 2002) and I-IX (generated by Dicer-mediated cleavage of ‛RevCen’ RNA; Djupedal *et al,* 2009; Figure 1B; Supplementary Table S2) and constructed DNA templates comprising the T7 promoter linked with the DNA sequences cognate to the siRNA sequences (Supplementary Table S4). The respective single stranded (ss)-siRNAs were generated by using T7 RNA polymerase (Supplementary Figure S3A). Of these, the siRNAs, denoted as ‛D-for’ and ‛E-for’, were radiolabeled and used to study the interaction with Swi6 (Supplementary Figure S3B, S3C) by Electrophoretic Mobility Shift Assay (EMSA; Figure 1C). Binding of Swi6 with the ‛D-For’ RNA was indicated by slow migrating bands I, II and III, representing different complexes (Figure 1C). The binding was specific, as it was competed out by excess of unlabeled D-For RNA (Figure 1C). No binding was detected with the complementary D-Rev RNA (not shown). Other ssRNAs: B, E and H (Figure 1C) and K (not shown), as well as I-IX; (Supplementary Figure S4A, S4B) also competed efficiently, suggesting the presence of conserved sequence elements in ‛A-K’ siRNAs (Figure 1C). The strength of binding/competition efficiency was in the order: B>D=E>H. A consensus Swi6/HP1-binding sequence obtained by sequence alignment was: G/C.A.G.T/C/A.A.G/T/A.G/C.G/T.G/C/A.T (Supplementary Figure S5). Interestingly, in contrast with the D-For RNA, a single band was observed for binding of Swi6 to the E-for RNA (Figure 1G, last lane) and the equilibrium binding constant, Kd, for Swi6/HP1 binding to ‛E-For’ RNA with a single binding site, was estimated to be 2.15±0.1μM (Figure 1E). [Likewise, analysis of binding isotherms for the binding sites I, II and III in the D-For RNA (Supplementary Figure S6) indicated that these are independent binding sites with Kds of 2.45µM and 1.79 µM for sites I and III, respectively, while that for site II was not considered as its isotherm did not show any dose-dependence (Supplementary Figure S6).]

It is worth noting that in the latter study EMSA was performed to detect binding of Swi6 to 100nt cenRNA and 150 and 750 nt GFP RNA in agarose gel and not acrylamide gel (Figure 3A in Keller *et al*, 2012). However, an EMSA using polyacrylamide gel failed to detect any binding of Swi6 to the ‘Cen100’ RNA, which was used in the mentioned study (Supplementary Figure 7A) or the ‛RevCen” RNA (Supplementary Figure 7B), the putative precursor of the heterochromatic si-RNAs (Djupedal *et al*, 2009). This difference may be due to the fact that EMSA in agarose gel can detect weaker binding due to larger pore size and less hindrance to the protein-RNA complex from migrating and dissociation. Consistent with that, we find that the binding affinity of Swi6 towards siRNA (this study) is ∼19-38-fold stronger than that reported for binding of Swi6 to a 20mer RNA based on NMR data (Kd=38±13µM; Keller *et al*, 2012). Thus, Swi6-HP1 shows a greater affinity and specificity towards the small RNAs like ‛D-For’ and ‘E-For’ than the longer euchromatic and heterochromatic precursor RNAs.

### Chromodomain-mediated binding of Swi6 to siRNAs

Earlier, the murine HP1α was shown to bind to RNA via its CD and this binding was influenced by the hinge domain, in particular, by the lysine triplet sequence in the hinge domain (Muchardt *et al*, 2002). Sequence comparison showed that this lysine triplet is conserved between the murine HP1 and Swi6/HP1 (residues 242-244; Figure 1F): mutation of the triplet lysine stretch abolished its RNA binding property (Muchardt et al., 2002). We found that, like HP1α, both the CD and CD-hinge but not the CSD region of Swi6 exhibit strong binding to the ‛E-For’ (used interchangeably with D-For RNA because of similar affinity; Figure 1G). Furthermore, the siRNA binding is drastically reduced in case of the Swi6^3K→3A^ protein (lysine triplet mutated to alanine; Figure 1D, right panel). Thus, unlike the binding of Swi6 to the *cen*RNA through the hinge domain (although detected only by EMSA in agarose gel), as reported by Keller *et al*. (2012), we find that, similar to mouse HP1α (Muchardt *et al*., 2002), using EMSA assay in acrylamide gel, that Swi6 binds to the *dg-dh* specific siRNA through the chromodomain and this binding is also regulated by the lysine triplet in the hinge domain.

These results show that different regions of Swi6 bind to the dg-dh specific siRNAs and to the cenRNA or GFP RNA (Keller et al., 2012) vs siRNAs: Swi6 binds to the *dg-dh* specific siRNAs through its chromodomain with 19-38-fold greater affinity and specificity than to the cenRNA, which is through its hinge domain.

### Swi6-HP1 is involved in protection but not generation of siRNA

We validated these results *in vivo* using RNA immunoprecipitation (RIP-seq) analysis. The radiolabeled RNA isolated from immunoprecipitates of extracts of cells expressing *GFP-swi6^+^*, *GFP-swi6^3K→3A^* or empty vector with anti-GFP antibody (Supplementary Figure S8A) was found to hybridize with blots containing DNA fragments of *dg*, *dh*, *cenH* (*dhk*) but not with a*ct1* (Supplementary Figure S8B). Denaturing urea/acrylamide gel electrophoresis of the radiolabeled size-fractionated siRNA indicated a qualitative and quantitative change in siRNAs in *swi6^3K→3A^* mutant, as compared with *swi6^+^* (Supplementary Figure S8A-S8C): while the Swi6-bound siRNAs show enrichment of the *dg-dh* sequences, the siRNAs bound to Swi6^3K→3A^ are depleted with respect to the *dg-dh* sequences and seem to bind smaller RNAs similar to *swi6Δ-*strain (Vector control, Supplementary Figure S8C). These results suggest that Swi6^+^ protein may bind to and protect the *dg*-*dh* siRNA from degradation but Swi6^3K→3A^ does not. RT-PCR analysis revealed the lack of bidirectional transcription in *swi6* strain as well as the transformed strains (Supplementary Figure S8D), while *dcr1*Δ strain showed accumulation of bidirectional transcripts specific for the *dg-dh* repeats, as shown earlier (Supplementary Figure S8D; Volpe *et al*, 2002). These results argue against role of Swi6 and Swi6^3K→3A^ in generation of siRNAs.

### Swi6-HP1 interacts with *dg*-*dh* specific siRNAs *in vivo*

Further, the size-fractionated small RNAs were subjected to RIP-Seq analysis. We observed no change in the genome-wide occupancy of siRNAs bound to Swi6p and Swi6^3K→3A^p (Figure 2A, left panel). However, we did observe a considerable depletion of siRNA corresponding to the *dg* and *dh* repeats in all three centromeres (Figure 2A, 2B; p= 0.001; Supplementary Figure S9B), the *dh* repeats in the *tlh1* and *tlh2* genes (Fig 2A and B; Hansen *et al*, 2006; Supplementary Figure S9B; p=0.02), and *cenH* region of the *mat* locus (Figure 2A, 2B) in the *swi6^3K→3A^* mutant as compared to *swi6^+^* strain. This is consistent with the report that Swi6 remains bound to sub-telomeric domains in the absence of telomeric repeats (Kanoh *et al*, 2005), implying prominent sub-telomeric heterochromatinization by Swi6 and associated siRNAs.

**Figure 2.**
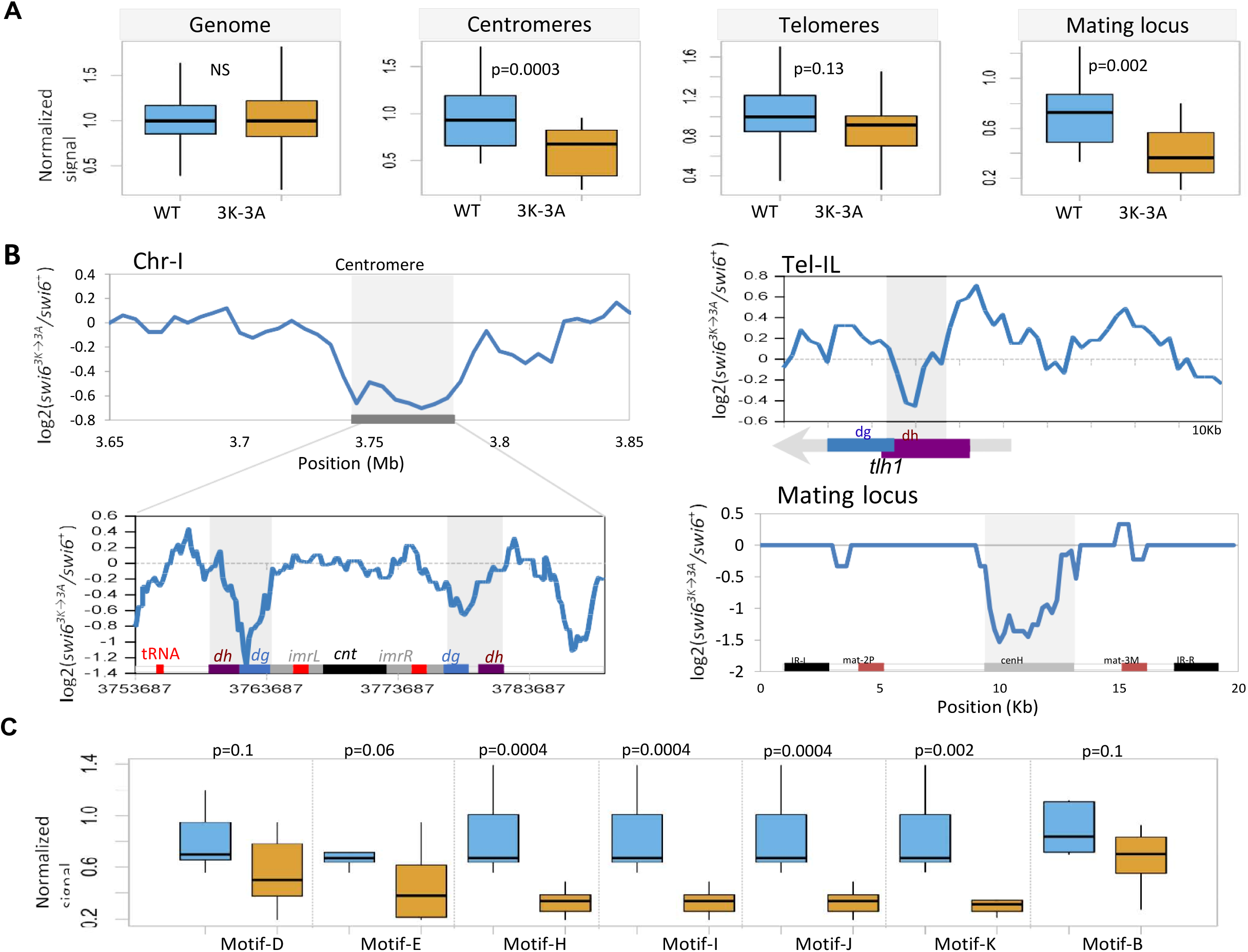
Localization of Swi6-bound siRNAs to *dg*-*dh* regions is abrogated in *swi6^3K→3A^* mutant. A Overall distribution of normalized small-RNA accumulation at centromeres, telomeres and mating-type locus in WT and *swi6^3K→3A^* mutant. B Chromosomal track of RIP-seq fold change (log2 *swi6^3K→3A^* /WT ratio) at and around centromere of chr-I at 5kb (top) and 200bp (bottom) resolutions, left telomere of chr-I (200bp resolution) and the mating locus (200bp resolution). Locus annotations are given at the bottom of the plot. Plots for other centromeres and telomeres are given in the Supplementary Fig10. C Distributions of normalized RIP-seq signals in *swi6^+^* and *swi6^3K→3A^* strains for regions containing the sequence motif(s) known to interact with Swi6. The p-values were calculated using two-tailed Mann-Whitney U tests.

To validate the EMSA results we analyzed the RIP-seq signals in the genomics bins containing the sequence motifs ‘A-K’ (Reinhardt & Bartel, 2002). The analysis clearly showed significantly reduced enrichment of the reported siRNAs ‛A-K’-bound to Swi6^3K→3A^ versus the Swi6^+^ protein (Figure 2C), confirming that these siRNAs do interact with Swi6 but not Swi6^3K→3A^ *in vivo*.

### Loss of siRNA binding abrogates heterochromatin localization of Swi6 and silencing

We further analyzed whether the *swi6*^3K→3A^ mutation affects the heterochromatin localization of GFP tagged Swi6. As reported earlier (Pidoux *et al*, 2000), ∼65% of *swi6^+^* cells contain 3 spots, ∼25% cells two spots and 10% cells one spot GFP-Swi6 (Supplementary Figure S10A, S10B). However, in *swi6^3K→3A^* mutant, only ∼20% cells each contained one, two or three spots or diffuse appearance (Supplementary Figure S10A, S10B), consistent with delocalization of Swi6^3K→3A^ from heterochromatin (Pidoux *et al*, 2000). We performed plate and ChIP assay to assess the effect of the *swi6^3K→3A^* mutation on silencing. Indeed, we observed derepression of the *ade6* reporter at the outer repeat *otr1R* and *ura4* at the inner repeat *imr1L* of *cenI* in *swi6^3K→3A^* mutant, as indicated by growth of pink colonies on adenine limiting media (Figure 3A, 3B) or enhanced growth of cells on plates lacking uracil, respectively (Figure 3E; Allshire *et al*, 1994). Complementing these results, ChIP assay showed reduced localization of Swi6 and Me2-K9-H3 at the otr1R::*ade6* (Figure 3C, 3D) and Swi6 at the *ura4* reporter, respectively in the *swi6^3K→3A^* mutant (Figure 3F, 3G). Similar loss of silencing at the *his3-telo* locus (Cooper *et al*, 1997) was indicated by enhanced growth of the *swi6^3K→3A^* strain on plates lacking histidine (Figure 3H, 3I), which is accompanied by reduced localization of mutant Swi6 at the *his3-telo* locus (Figure 3J, 3K). The abrogation 8of silencing of the *mat2*-linked *ura4* reporter in the *swi6^3K→3A^* mutant was indicated by lack of restoration of FOA sensitivity in *swi6Δ* strain by the *swi6^3K→3A^* gene (Supplementary Figure S10C, S10D). Furthermore, in the homothallic, efficiently switching background (*h^90^*), the *swi6^+^* gene restored efficient switching to the *h^90^ swi6Δ* mutant, as indicated by enhanced iodine staining of the transformants (Moreno *et al*, 1991; Materials and Methods), but the mutant *swi6^3K→3A^* did not (Supplementary Figure S10E). Thus, the siRNA-binding property of Swi6 is important for silencing and heterochromatin localization of Swi6 and H3-K9-Me2 and mating type switching.

**Figure 3.**
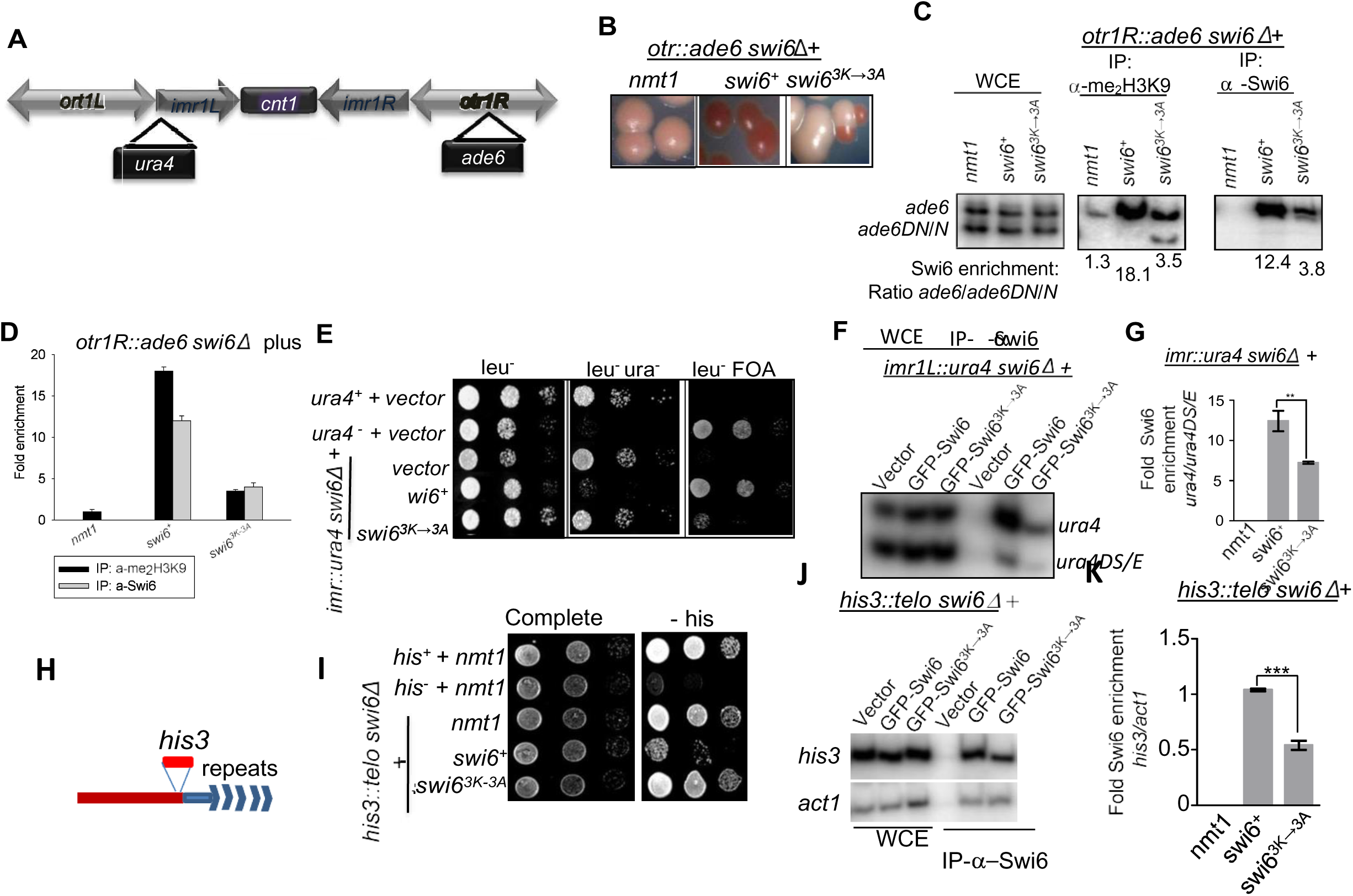
Abrogation of silencing in *swi6^3K→3A^* mutant accompanies Swi6 delocalization at the outer repeat *otr1R* and *imr1L* regions of *cenI* and *his3-telo*. A Diagrammatic representation of the genotype of the strain. Centromere of each fission yeast chromosome comprises of a central core (*cnt*), immediately flanked on both sides by inverted repeats (*imr*); *imr* on either side are flanked by outer repeat regions (*otr*). Reporter genes *ura4, ade6* and *his3* have been inserted into the *imr*, *otr* of centromere and telomeric repeats, respectively of chrl. The strain has a *swi6Δ* mutation and carries internal deletions in native *ura4* locus, *denoted as ura4DS/E,* and in *ade6*, denoted as *ade6DN/N*, which act as euchromatic controls in ChlP assay. B-D Derepression of the *otrIR::ade6* locus in *swi6^3K→3A^* mutant, showing pink/white colonies on adenine limiting plates (B). C ChIP assay showing reduction of Me2-K9-H3 and Swi6 at the *ade6* locus in *swi6^3K→3A^* mutant. Ratio of the heterochromatic *ade6* and euchromatic *ade6DN/N* locus is shown D Quantitation of data shown in (C). E Spotting assay showing inability of *swi6^3K→3A^* to restore silencing at the *imrl::ura4* locus in *swi6Δ* mutant, as indicated by growth on plates lacking uracil and lack of growth on FOA plates. *F* ChIP assay showing delocalization of Swi6 from the *imrl::ura4* locus in the *swi6*^3K→3A^ mutant, where *ura4* represents the heterochromatin and *ura4DS/E* the euchromatin. G Quantitation of the data showing enrichment of Swi6 at *ura4* versus *ura4DS/E* in (F).H-K Derepression of the subtelomeric*his3* locus in *swi6*^3K→3A^ mutant. H gives a schematic representation of the *his3* gene inserted distal to the telomeric repeats, denoted as *his3*-*telo*. I Spotting assay showing the enhanced growth of the strain having *his3-telo* reporter in presence of the *swi6* mutation on plates lacking histidine. J ChIP assay reduced localization of mutant Swi6^3K→3A^ at the *his3-telo* locus as compared to wt. *act1* was used as a control. K Quantitation of data shown in (J).

### Swi6-HP1 exhibits strong binding to siRNA-DNA hybrids *in vitro* and *in vivo*

We envisaged that the si-RNA binding by Swi6/HP1 could facilitate its recruitment to the cognate complementary sequences in the genome. We speculated that Swi6^+^ may also bind to the siRNAs as RNA-DNA hybrid. Indeed, EMSA experiment confirmed the presence of three binding sites, named I, II and III, for Swi6 on the ‛D-For’ RNA-DNA hybrid with very high affinity with Kd of 0.08 μM, 0.09 μM and 0.73 μM, respectively (Figure 4A, 4B), while Swi6^3K→3A^ showed no binding (Figure 4A, right panel).

**Figure 4.**
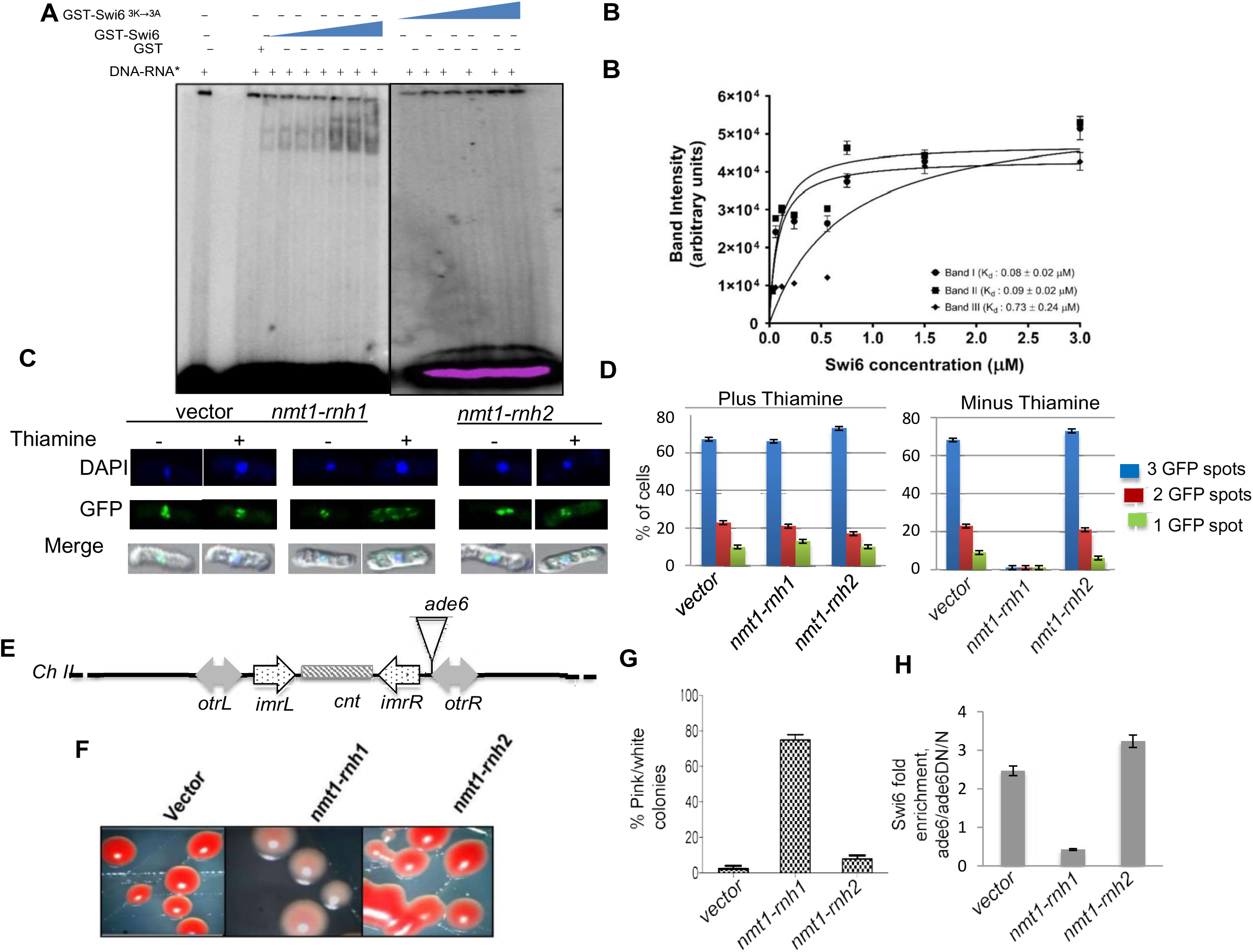
Role of Swi6 binding to the *dg*-*dh* sequences as RNA-DNA hybrids in silencing. A Swi6 shows strong affinity towards the RNA-DNA hybrid. EMSA assay was performed using the hybrid between the D-For RNA and its complementary DNA sequence and purified GST-Swi6 or GST-Swi6^3K→3A^. GST was used as control. B Data in (A) was plotted to determine the equilibrium binding constant, Kd. C Delocalization of GFP-Swi6 upon overexpression of *rnh1* but not *rnh201*. Fluorescence micrographs showing the effect of overexpressed *nmt1*-*rnh1* and *nmt1*-*rnh201* on the subcellular localization of Swi6 with respect to the three heterochromatic loci, *mat*, *cen* and *tel* as distinct green foci. The strain had the native copy of *swi6* gene chromosomally tagged with GFP. Nuclei were stained with DAPI. Strains expressing GFP-tagged Swi6 was transformed with vector alone, *rnh1* or *rnh201* under the control of *nmt1* promoter. Transformed cells were grown in the presence (+) or absence (-) of the repressor, thiamine, visualized under confocal microscope and photographed. D Microscopic data shown in (C) was quantified as distinct green foci pertaining to GFP-Swi6 subcellular localizing to the heterochromatic loci. Quantitation of the localization of GFP-Swi6 as three, two or one spots. E-H Overexpression of *rnh1* causes loss of silencing at the *otr1R::ade6* locus. E Schematic representation of the centromere I having an *ade6* reporter gene inserted into the *otr1R*. F The effect of expression of empty vector alone (*nmt1*), *nmt1*-GFP-*swi6* or *nmt1*-GFP- *swi6^3K→3A^* on the expression of the *ade6* reporter. Colors of the colonies indicated the state of the *ade6*reporter. Red colonies indicate suppression of *ade6*repression while pint/white colonies indicated repression. The data are represented in terms of percent white colonies. G Plot showing the per cent colonies giving pink/white appearance in different transformants, as shown in (F). H Delocalization of Swi6 from the *otr1R::ade6* locus in the transformants shown in (F). Derepression of the *ade6* reporter at *otr1R* repeat as a result of overexpression of *rnh1* and *rnh201*, as observed in (G) was biochemically verified through Swi6 enrichment at the locus through ChIP assay.

To confirm the RNA-DNA binding by Swi6/HP1 and its role in heterochromatin formation *in vivo*, we checked the susceptibility of heterochromatin formation to RNase H, which cleaves the RNA-DNA hybrids. Indeed, overexpression of *rnh1* (RNase H1) but not *rnh201* (RNase H2 subunit A) in cells expressing GFP-tagged Swi6 caused delocalization of GFP-Swi6 as visualized by confocal microscopy (Figure 4C, 4D; Supplementary Figure 11). This was accompanied by loss of silencing of the *ade6* reporter at the *otr1R* repeat of *cenI* and *mat3* locus, as indicated by growth of pink colonies on adenine limiting medium (Figure 4E-4G and Supplementary Figure 12A-12C), while ChIP assay showed reduced localization of Swi6 at both loci (Figure 4H; Supplementary Figure S12D) in cells transformed with *rnh1* but not by empty vector or *rnh201*. Overexpression of *rnh1* also elicited loss of silencing at the *mat2P* locus in a strain containing a stable *mat1M* locus, as indicated by iodine staining assay (Moreno *et al*, 1991; Materials and Methods; Supplementary Figure S12E-S12G) and ChIP assay showed loss of Swi6 localization at the *mat2*-linked *ura4* reporter (Supplementary Figure S12H). These results confirm that localization of Swi6 to the *dg*-*dh* repeats (and spreading, as in case of *mat2::ura4* and *mat3:: ade6*) is dependent on its binding to the cognate RNA-DNA hybrid.

To check whether Swi6/HP1 binds and protects the *dg-dh* repeats existing as RNA-DNA hybrid *in vivo*, we performed the DNA-RNA Immunoprecipitation (DRIP) experiment (Ginno *et al*, 2012) using the monoclonal antibody against RNA-DNA hybrid (S9.6; ATCC HB-8730). Chromatin samples that were immunoprecipitated with the antibody were treated with RNaseH and the RNaseH-resistant *dh* regions were quantitated by real time PCR (light grey bars; Figure 5A). Treatment with RNaseH *in vitro* caused a 2-fold reduction of the *dh* signal in *swi6^3K→3A^* as compared to the *swi6^+^* cells and also in *swi6* cells transformed with the empty vector and plasmid expressing *swi6^3K→3A^*, as compared to plasmid expressing *swi6^+^* gene (Figure 5A), supporting a role of Swi6 in binding and protecting the RNA-DNA hybrid.

**Figure 5.**
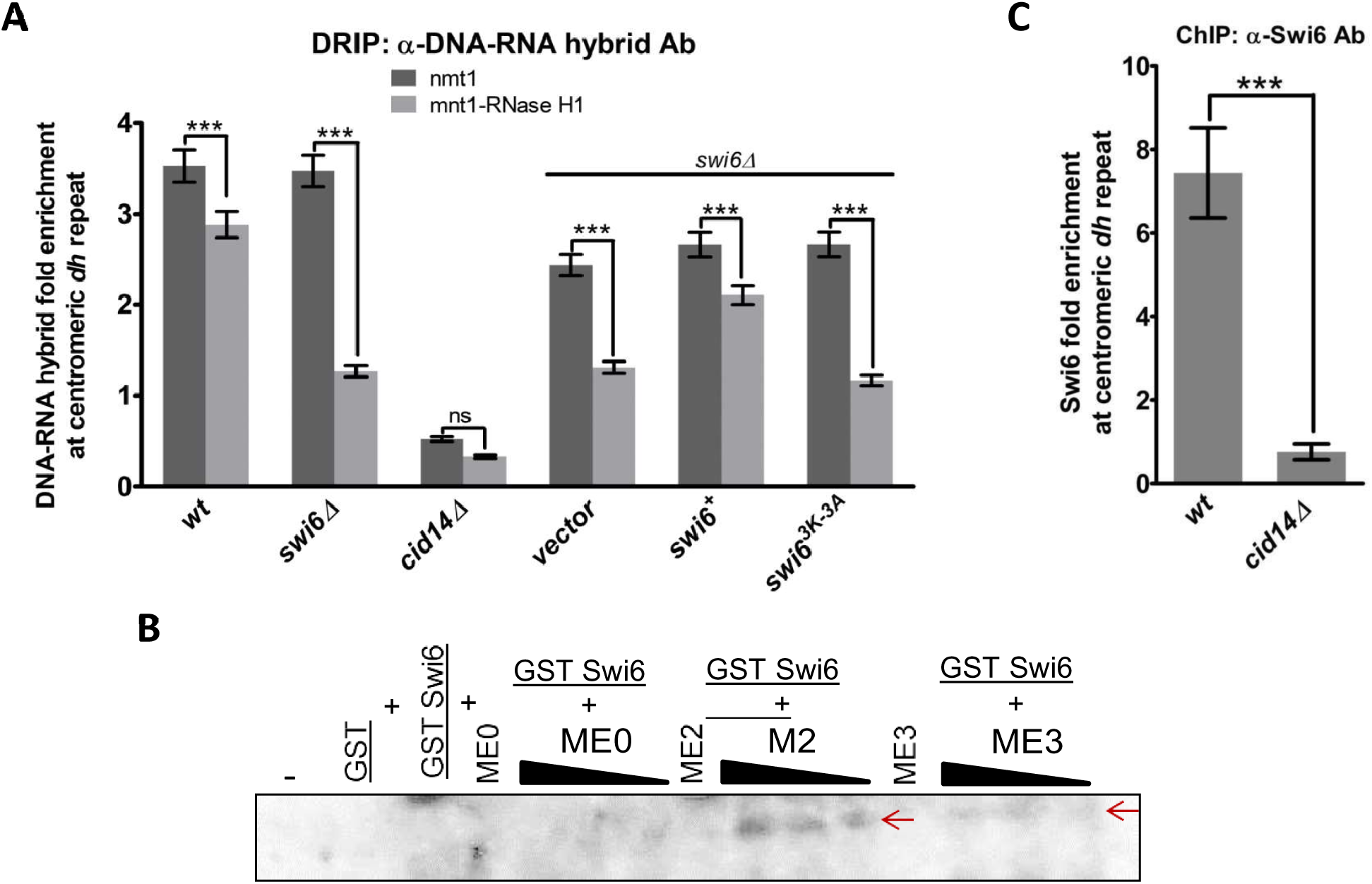
Swi6 binds to and protects the RNA-DNA hybrid at the *dh* region *in vitro*. A ChIP assay was performed to quantitate the *dh* regions enriched with anti-RNA-DNA hybrid antibody in the indicated strains. B The RNA-DNA hybrid shows preferential binding to the Me2-K9-H3. EMSA was performed to detect binding of Swi6 to the RNA-DNA hybrid (D-For RNA/DNA duplex) in presence of unmethylated (M0), Lys9 dimethylated (M2) and Lys9 trimethylated H3 (M3) peptides. No binding of the hybrid was observed with any of the peptides. To be able to observe the supershift (indicated by arrowheads) in presence of the peptides, electrophoresis had to be performed longer to run out the labeled hybrid, which also led to a diffuse signal. C si RNA generation is required for Swi6 enrichment at *dh* region. ChIP assay was performed to quantitate the *dh* region associated with Swi6 in wt versus *cid14Δ* mutant.

### siRNA-DNA-Swi6 complex binds specifically to Me2-K9-H3

Earlier, using SPR, the GFP 20-mer RNA was shown to show preferential binding with Me2/Me3/-K9-H3 over Swi6, especially at above-stoichiometric ratios of RNA to Swi6 (Figure 6E; S6D in Keller *et al*, 2012). Furthermore, we performed EMSA assay to determine whether the binding of Swi6 to the D-For RNA/DNA hybrid affected its binding to the Me2/Me3-K9-H3 peptide. Surprisingly, we observed a dose-dependent super-shift in the binding of Swi6 in presence of Me2-K9-H3 (Figure 5B, M2) and to a lesser extent with Me3-K9-H3 peptide (Figure 5B, M3) but not with the unmethylated H3 peptide (Figure 5B, M0).(Owing to drastically slower mobility of the peptide bound complexes, electrophoresis was carried out for longer period due to which the bands for the RNA-Swi6 bound complexes were allowed to run out of the gel). Thus, the high affinity binding of Swi6 to the *dg*-*dh* specific RNA/DNA hybrid facilitates the binding to Me2-K9-H3 rather than competing against it.

**Figure 6.**
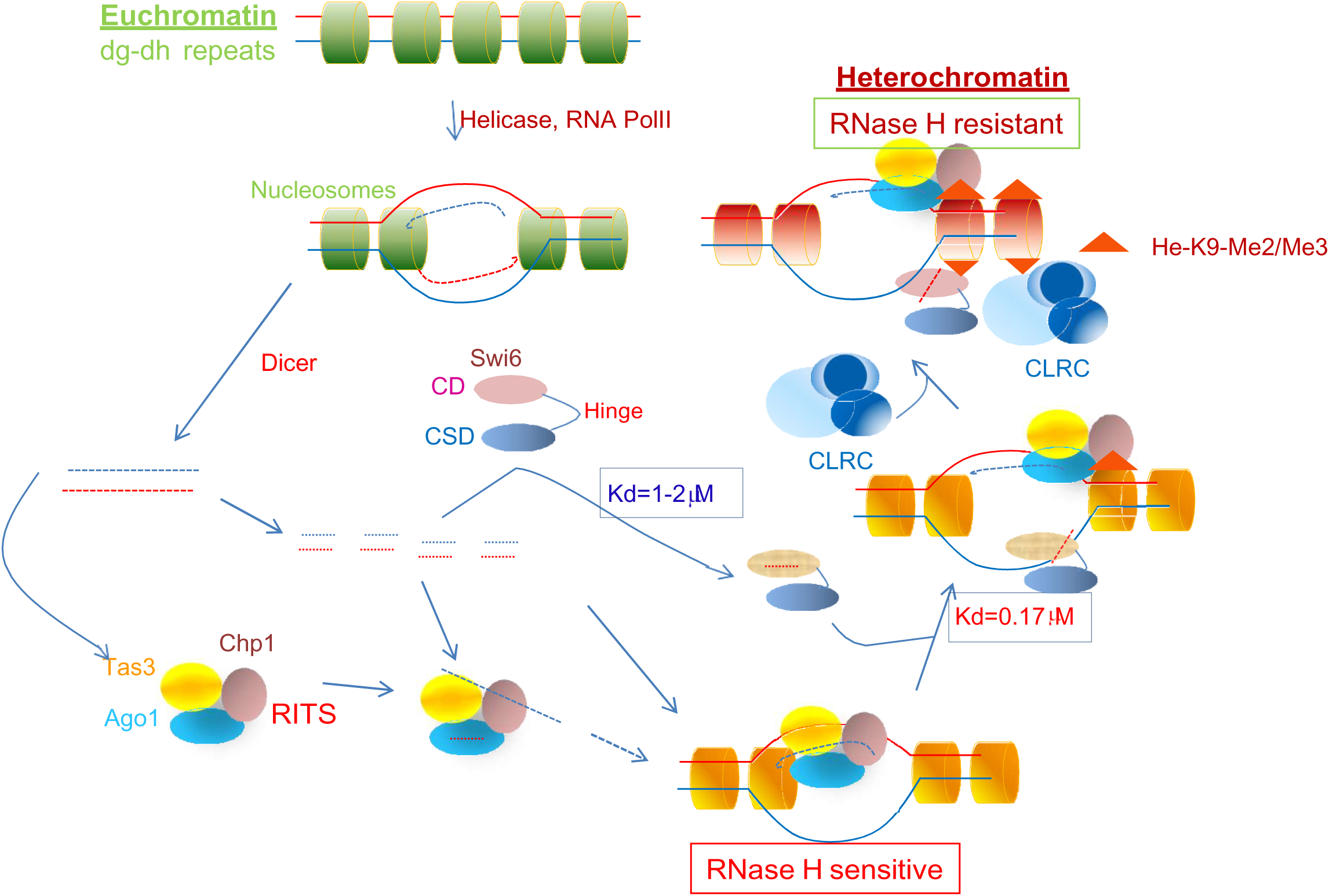
Speculative model about the role of nucleic acid binding function of Swi6/HP1 in initiating RNAi dependent heterochromatin assembly in Fission Yeast. Based on data described here, Swi6/HP1 binds with moderate affinity to the single stranded siRNA mapping to the dg-dh repeats. This helps it to be recruited to chromatin at the stage in the RNAi pathway where the RITS complex is already tethered through the nc transcript. This stage is sensitive to RNase H. The recruitment of Swi6-ss-siRNA complex is facilitated by the occurrence of the complementary DNA strand in single stranded form. The high affinity binding of Swi6 to the RNA-DNA hybrid helps to retain Swi6 and makes its localization RNase H-resistant. Subsequent recruitment of the CLRC complex helps to methylate the histones and promote formation of a more stable ternary complex with Swi6-HP1.

### Heterochromatin recruitment of Swi6-HP1 depends on siRNA generation

It was shown earlier that the level of siRNAs originating from the outer repeats of centromere is reduced in the *cid14Δ* mutant, which is defective in polyadenylation of RNAs and their subsequent degradation by the exosome pathway, resulting in accumulation of pre-mRNAs and ribosomal RNAs (Wang *et al*, 2008). As the siRNA level is reduced in *cidl4* mutant, we speculate that siRNA binding may be the first step in recruitment of Swi6 to the *dg*-*dh* repeats (Muchardt et al., 2002). This is in agreement with the reported dependence of Swi6-HP1 localization to centromeres on Dicer (Woolcock *et al*, 2011). In line with previous results, a reduction in the level of siRNAs corresponding to the dg-dh repeats should result in decrease in Swi6 localization to the cognate sites. Indeed, results of real time PCR confirm that the level of Swi6 recruitment to the *dh* repeats is reduced in the *cid14Δ* mutant (Figure 5C).

### Mutant Swi6^3K→3A^ shows shift in binding to euchromatic transcripts

Keeping in mind the reported binding of Swi6 to the cen and GFP RNAs, albeit at much lower affinity, we asked whether the loss of binding by Swi6^3K→3A^ mutant protein to the siRNAs generated from the *dg-dh* repeats may render the unbound protein available for binding to non-heterochromatic RNAs. We compared the regions with >2-fold increase in siRNA enrichment in mutant with the regions exhibiting >2-fold increase in WT, for the enrichment of Swi6, RNA PolII, Me3-K36-H3, from the ChIP-seq signals and the gene expression values. As expected, siRNAs bound to the Swi6^3K→3A^ protein were found to map to regions that show less association with Swi6^+^ (Supplementary Figure S13). Interestingly, mutant-bound siRNAs mapped to regions showing greater enrichment of RNA PolII and Me3-K36-H3, which are known to be associated with transcription elongation (Morris *et al*, 2005), as well as higher gene expression values, implying the loss of specificity of Swi6^3K→3A^ for *dg*-*dh* siRNAs and a general non-specific shift towards abundantly transcribing regions (Supplementary Figure S13). Most interestingly, a similar shift occurs in case of the ‛Cen100’ RNA (SPAC15E1.04); while no significant enrichment was observed in Swi6+ cells, there was ∼70% increase in Swi6^3K→3A^ Cells (Supplementary Figure S14).

## Discussion

This study was undertaken to address the causal connection between the RNAi and heterochromatin pathways in fission yeast. It was shown earlier that *rnai* mutants undergo loss of heterochromatin structure and silencing due to depletion of histone H3-Lys-9 dimethylation and Swi6/HP1 at the heterochromatin loci (Volpe *et al*, 2002). However, despite discovery of various complexes, like ARC, RITS, RDRC, etc, no direct connection between the two pathways has been forthcoming. An indirect involvement of Chp1 as a component of the RITS complex has been suggested to play a role in recruitment of Clr4-containing CLRC complex through Stc1 (Bayne et al., 2010); however, an important caveat against this proposal is that Chp1 recruitment, in turn, is dependent on a prior H3-Lys9 methylation by Clr4.

An earlier report had shown a role of binding by mouse HP1 to RNAs in determining its heterochromatin localization in mammalian cells (Muchardt *et al*, 2002). However, a similar role of RNA binding and heterochromatin localization by Swi6/HP1has not yet been reported in fission yeast. Rather, Swi6/HP1 was shown to bind general RNAs with low binding affinity, which competes against the binding of Swi6 to H3-Lys9-Me2/Me3, specially at above-stoichiometric RNA/Swi6 ratio (Keller *et al*, 2012): accordingly, it was proposed that RNA may sequester Swi6 away from methylated H3 and get channeled to the exosome pathway.

Here, we have investigated whether, like the mammalian HP1 (Muchardt *et al*, 2002), Swi6 binds to specific siRNA sequences (that are generated by the RNAi-mediated cleavage of the siRNA precursor RNAs, which are transcribed from the centromeric repeats) or to the precursor transcripts. Our results show that **i**) indeed Swi6 shows high binding affinity and specificity towards the peri-centromeric siRNAs (Reinhardt & Bartel, 2002; Djupedal *et al*, 2009), with an increasing order of affinity towards single-stranded siRNAs (Kd=1-2 μM) and siRNA-DNA hybrid (Kd=30nM-0.7 μM). These binding affinities are, respectively, >30 to 1300-fold stronger than that reported for binding to the 20-mer RNA (Kd=38 μM; Keller *et al*, 2012). Notably, in contrast with the earlier work showing binding of Swi6 to the ‘Cen100’ or GFP RNA by EMSA assay in agarose gel, the EMSA assay in acrylamide gels (present study; Supplementary Figure S7) showed no binding of Swi6 to either the ‘Cen100’ RNA, or the ‛RevCen’ RNA, from which the siRNAs denoted as VI-IX (used in this study) are derived (Djupedal *et al*, 2009), indicating a preferential binding by Swi6 to the siRNAs instead of the siRNA precursors. **ii**) This binding to siRNA is ascribed to the CD although, surprisingly it is influenced by the hinge domain. In contrast, Swi6 was shown earlier to bind to the GFP or Cen100 RNA through the hinge domain and not the CD (Keller *et al*, 2012), indicating a fundamental difference in the nature of binding of Swi6 to GFP/Cen100 RNA versus siRNAs. **iii**) The *in vitro* binding of Swi6/HP1 to the peri-centromeric (*dg-dh* specific) siRNAs is validated *in vivo* by results showing enrichment of *dg-dh* specific siRNA species bound to Swi6 and depletion of their levels bound to the Swi6^3K→3A^ mutant protein. **iv**) Localization of Swi6 to heterochromatin is abolished in the triplet lysine mutant *swi6^3K→3A^*, and in *swi6^+^* cells treated with RNase H1. This localization is likely to be causally linked to Swi6 binding to and stabilizing the cognate siRNA-DNA hybrid. **v**) In support, reduction in the level of siRNA in *cid14Δ* mutant, which is defective in polyadenylation of mRNAs and their subsequent degradation by exosome, is also accompanied by reduction in heterochromatin localization of Swi6. **vi**) There is a shift from Swi6^+^ binding to heterochromatin-specific siRNAs to euchromatin-specific RNA by the Swi6^3K→3A^ mutant protein.

An important feature of the binding of Swi6 to siRNA is that it promotes the binding to the H3-K9-me2 peptide (Figure 5B), which is in contrast to the ability of GFP RNA to compete out the binding of Swi6 to the H3-Lys9-Me2 (Keller *et al*, 2012). This distinction can be ascribed to the fact that Swi6 binds to siRNA primarily through the chromodomain (CD, present study), while the binding of Swi6 to the Cen100 RNA is reported to occur through the hinge domain (Keller *et al*, 2012). Thus, Swi6 may interact by two distinct modes to RNA: a high affinity binding of *dg-dh*-specific siRNA to the CD, which is distinct from but supports the subsequent binding of H3-Lys9-Me2 and a low affinity site in the hinge domain for binding to non-coding Cen/mRNAs, that, paradoxically competes out the binding to the H3-Lys9-Me2 peptide. The molecular basis of this paradoxical result remains to be addressed.

To address this apparent discrepancy, we envisage a dual function of Swi6. Under normal conditions, Swi6 can bind with high affinity to the siRNAs generated by Dicer-mediated cleavage of the RNA precursors transcribed by RNA PolII from the centromeric *dg-dh* repeats. An integral role of siRNAs in heterochromatin assembly is also supported by the finding that only the siRNA-rich (and not siRNA void) sub-domains of dg and dh regions could nucleate heterochromatin at ectopic sites (Buscaino et al., 2013). By virtue of the enhanced binding of the siRNA-Swi6 complex can get sequestered to the cognate DNA sequences in the *dg-dh* repeats as RNA-DNA hybrids independently of H3-Lys9-Me2; this binding to siRNA-DNA hybrid could help to lock the complex in the chromatin and recruit Suv39/Clr4 H3-Lys9 methyltransferase to initiate heterochromatin formation, as suggested earlier (Haldar *et al*, 2011; Al-Sady et al, 2013). It is possible that apart from normal RNA, cells also produce a large amount of incomplete transcripts or those containing termination codons, especially under stress conditions (Buhler et al., 2008); these excess RNAs may sequester Swi6 from the heterochromatin regions and get channeled to the exosome pathway. It is possible that the higher amount of these RNAs may overcome the lower affinity and bind to Swi6. In fact, this scenario resembles the H3-K9-Me binding mode of Swi6 at below stoichiometric ratios of RNA to Swi6 (Keller et al., 2012) is supported by the result showing that the Swi6^3K→3A^ mutant protein binds more strongly to the Cen100 than the wt Swi6 protein (Supplementary Figure S14). These considerations point to a dynamic role of Swi6 in RNA surveillance, physiology and turnover.

A moot point considering the earlier report of Keller *et al* (2012) is whether Swi6 shows binding to the siRNA or to the precursor RNAs/mRNAs. In this regard, we find that while Swi6 does not show any detectable binding to the ‛RevCen’ RNA, it does exhibit strong binding to the siRNAs denoted as ‛VI’ to ‛IX’, which are generated by Dicer cleavage of the ‘RevCen’ RNA (Supplementary Figure S4; Djupedal *et al*, 2009). This difference may be ascribed to the possible sequestration of the regions of the ‘RevCen’ RNA corresponding to the sequences ‛VI’–‛IX’ cither in the double-stranded form and/or in a protein-bound state. According to this scenario, the siRNAs (e.g. VI-IX) are more likely to exist in single-stranded form and thus be more accessible for binding to Swi6. In contrast, the ‛RevCen’ RNA and ‛Cen100’, a representative mRNA molecule, may bind with lower affinity due to its partial double-stranded nature, an inference supported by lack of detectable binding as assessed by EMSA assay in acrylamide gel. (Binding in agarose gel but lack of binding in polyacrylamide gel may also reflect the comparatively lower stability of the Swi6-Cen100 or Swi6-GFP RNA complexes, which may lead to their dissociation in acrylamide gel but not in agarose gel; Supplementary Figure S7). Thus, Swi6 may bind to siRNAs engendered by cleavage of the precursor RNAs transcribed from the centromeric repeats, in a sequence-specific manner at moderately high affinity (Kd∼1-2 μM), which is ∼20-40-fold greater than that reported for the ‛Cen100’ ‛RNA’ (38µM; Keller *et al*, 2012). It also appears that the Cen100 RNA, and possibly RevCen RNA, could bind more strongly to the Swi6^3K→3A^ mutant protein, possibly through hingedomain but not the CD, as argued above.

To get an idea about the sequence specificity we used the MEME suite (Bailey *et al*, 2009). We compared the siRNA sequences showing the highest affinity to Swi6, i.e., E, B, K (Reinhardt & Bartel., 2002) and I, IV, VII, I (Djupedal *et al*, 2009) and arrived at a consensus sequence G/C.**A**.**G**.T/C/A.**A.G**/T/A.G/C.G/T.G/C/A.T (Supplementary Figure S5). We did not include the sequence D as it seems to contain at least 2 binding sites. A comparison of the consensus with the sequence D showed that a subset sequence AG does occur twice in “D’ sequence. It would be interesting to map the actual binding residues by experiments involving RNA foot-printing combined with mutagenesis. Most interestingly, Swi6 shows extremely high affinity to the siRNA-DNA hybrid with Kd of up to 30nM (Supplementary Fig. 15). Further work is needed to decipher the sequence specificity in Swi6 binding to RNA and testing its role in heterochromatin assembly.

Based on these results we propose the model that Swi6 may be initially bound to the siRNAs generated by RNAi-mediated cleavage of the centromeric transcribed RNAs. A still stronger binding to the respective sequences in the form of RNA-DNA hybrids may help to lock the Swi6 to those sites in the centromere. This type of binding is different from that of Swi6 to Cen100 in that i) it is 19-38 stronger and ii) it does not displace the binding of Swi6 to H3-Lys9-Me2; rather the binding of Swi6 to the RNA-DNA hybrid facilitates the binding to the H3-Lys9-Me2. Thus, the nature and site of binding of Swi6 to siRNA and siRNA-DNA hybrid seems to be different from the one to H3-Lys9-Me2 unlike that of Cen100 (Keller *et al*, 2012; Supplementary Figure S7); while the binding of Swi6 to Cen100 competes for the binding by H3-Lys9-Me2, its binding to siRNA is independent of but synergistic to that of H3-Lys9-Me2.

Here it is pertinent to compare and contrast the binding characteristics of CD of Chp1, Swi6 and Clr4. Similar to our results, an earlier study also showed lack of binding of Swi6 to the siRNA precursor, similar to the ‛RevCen’ RNA (Figure 7B in Ishida *et al*, 2012). However, a similar RNA was shown to bind to Chp1 and this binding was further enhanced by binding to H3-Lys9-me2. (Ishida *et al*, 2012). This study also ascribed the RNA binding by Chp1 to the chromodomain (CD) region and, more specifically, to the basic residues in the C-terminal α-helical region of the CD. Furthermore, it was shown that CD of Clr4 could not bind to RNA on its own but did so in presence of the H3-Lys9-Me2. A comparison of the CD sequence of Chp1, Clr4 and Swi6 reveals a greater preponderance of basic residues in the C-terminal α-helical region in the CD of Chp1, while in case of Swi6 the basic residues are more prevalent in the N-terminal β1 region of the CD (Figure 4D in Ishida *et al*, 2012). These sequence differences may be responsible for diversity insequence specificity among the CD proteins.

Another pertinent issue is the relative affinity of the CDs towards H3-Lys9-Me2 and nucleic acids. CDs of Chp1, Clr4 and Swi6 exhibit decreasing order of affinity towards H3-Lys9-Me2, with Kd of 0.55μM, 2.86μM and 10.28μM, respectively. Thus, Swi6 binds with ∼5-10-fold greater affinity to to siRNA (Kd ∼1-2μM- this study) than to H3-Lys9-me2 (Kd of 10.28μM, Schalch *et al*, 2009) and in particular, to the RNA-DNA hybrids (30-700 fold; see Supplementary Table S6). In this regard, Schalch *et al* (2009) made a puzzling observation: although the mutants of Chp1 with reduced binding affinity to H3-Lys9-Me2 caused its defective localization away from heterochromatin, heterochromatin structure was still maintained, which suggested some overlapping mechanism in heterochromatin formation independent of H3-K9-Me-binding. Further, it is also puzzling how an ∼4 fold stronger binding of Swi6 to H3-K9-Me2 (Kd= 10.28μM) than to GFP20 mer RNA (Kd= 38μM) can explain the displacement of Swi6 from H3-K9-Me2/3 by RNA, except perhaps under condition of of excess RNA. Considering these points, we speculate that the chromodomains of Chp1, Clr4 and Swi6 may perform a division of labor through differentiation of the role of their CD regions by binding to either the precursors RNA or its siRNA derivatives, for achieving diverse outcomes.

Another point needs mention here. It was shown earlier that *cid14Δ* mutant shows a reduced Amount of heterochromatic GFP mRNA and a ∼ 2-fold reduction in the level of H3-K9-Me2 at mat3::GFP (with p= 0.2; Figure 2C, Keller et al., 2012) and tlh1/2 (p=0.28; Figure 2D, Keller et al, 2012). In contrast, a greater reduction in the level of H3-K9-Me2 was observed in *clr4Δ* mutant as compared to wild type at mat3M::GFP locus (p=0.01; Figure 2C, Keller et al., 2012) and at tlh1 (p=0.3 x10^-8^; Figure 2D; Keller et al., 2012). On the other hand, we show a complete loss of Swi6 at the dh repeats (p= <0.0001, Figure 5C, this study)

A rather puzzling observation is the differences in the effects of RNaseH1 and RNaseH2 on silencing. Studies have shown that while both the enzymes degrade the RNA in the RNA-DNA hybrids at sites similar to the Okazaki fragments, there are some subtle differences in their mechanism and site of action. RNase H1 and RNase H2 are differentially regulated to process RNA-DNA hybrids. RNase H1 can function at Okazaki fragments during S phase, where it can interact with RPA and localize constitutively to the stable R-loops via RPA-coated displaced ssDNA. Interaction with RPA further enhances the activity of RNseH1 (Nguyen *et al*, 2017; Petzold *et al*, 2015). It removes R loops but does not induce unwanted nicks in DNA at rNMPs. In contrast, RNaseH2 is a housekeeping enzyme that deals with majority of RNA-DNA hybrids (rNMPs and R-loops) at post-replication stage (Zimmer & Koshland, 2016). Thus, it is possible that the RNA-DNA hybrids may be susceptible to cleavage only during S phase. A trivial possibility is that RNaseH2 is composed of three subunits rnh201, rnh202 and rnh203; since only rnh201 was overexpressed alone, it may not exert any effect.

Summing up, we report that Swi6/HP1 displays a hierarchy of binding affinities to RNA and DNA sequences: i) weak affinity to the euchromatic transcripts like ‛Cen100’ or GFP 20-mer RNA, as reported earlier (Keller et al., 2012), ii) moderately high affinity with sequence specificity towards the ss- siRNAs corresponding to the *dg-dh* repeats, which is >15-fold stronger than that towards ‘Cen100’ and iii) strongest binding (over 220-fold) to the *dg*-*dh* sequences as RNA-DNA hybrid. This property of Swi6/HP1 may predominantly govern its specific localization at centromeric and centromere-like repeats, in a *cis*-acting manner, to the cognate sites in DNA and helps to initiate heterochromatin formation and silencing. This mechanism of Swi6/HP1 recruitment is independent of histone methylation but dependent on RNAi pathway. We speculate that the siRNAs bound to Swi6/HP1 may be chaperoned to the *dg*-*dh* sequences by sequence complementarity. The cognate sites may be unwound due to transcription by RNA PolII and/or Rdp1 and/or association with the RITS complex: these sites also exist as an RNA-DNA hybrid and are susceptible to RNase H1 (Shimada *et al*, 2016). This window of opportunity may allow Swi6 trafficking from the RNA-bound to the RNA-DNA-bound form, facilitated by the stronger binding of Swi6. The stronger binding may help to lock Swi6 stably to RNA-DNA hybrid rendering it resistant to RNase H. This model of Swi6 recruitment playing a role in heterochromatin at centromere dg-dh sequences is supported by the finding that these regions are enriched in fractions enriched immunoprecipitated with anti-RNA-DNA hybrid antibody (Dutrow et al., 2008) and RNAi-dependent heterochromatin formation is dependent on formation of DNA-RNA hybrid (Nakama et al., 2012). Furthermore, using the dam-ID technique, Swi6 and Dcr1 were shown to be localized to *cenII, mat* and tel: importantly, Swi6 localization was dependent on Dcr1 (Woolcock et al., 2011).

Subsequently, the Swi6-bound Clr4 may catalyze lysine methylation, as proposed earlier (Haldar *et al*, 2011; Al-Sady *et al*, 2013) and further heterochromatin spreading may occur by binding of Swi6/HP1 to Me2-K9-H3 (Figure 6). This scenario can explain the apparent discrepancy between the delocalization of Swi6 upon overexpression of RNaseh1 and protection of RNA-DNA hybrids by Swi6. Independent support comes from the enrichment of the *dg*-*dh* and *cenH* regions as RNA-DNA hybrids (Dutrow *et al*, 2008). In contrast, the Swi6^3K→3A^ protein’s binding sites may shift from the *dg-dh* repeats to the euchromatin regions because of low-affinity, promiscuous RNA binding property, as is indeed observed (Supplementary Figure S14).

We propose that because of evolutionary conservation, the metazoan orthologs of Swi6/HP1 may facilitate the heterochromatinization of centromeric repeats by a similar mechanism. Interestingly, the chromo- and hinge domains of Swi6/HP1 are conserved between *S. pombe* and higher eukaryotes and are interchangeable (Wang et al., 2000). Furthermore, the centromeres in higher eukaryotes also contain repeat sequences. Thus, the variations in the sequences in these domains in the metazoan orthologs of Swi6-HP1 may allow for their species-specific binding to the centromeric repeat-specific siRNAs and thereby regulate their binding to the cognate sites in the respective genomes as RNA-DNA hybrids in a sequence-specific manner. Given the conservation of components of RNAi and heterochromatin between the fission yeast and higher eukaryotes, the findings of this study should serve as a paradigm for understanding the connection between RNAi and heterochromatin assembly in eukaryotes.

## Materials and methods

### Strains and plasmids

The strains used in this study are listed in Supplementary Table4.

The *rnh1* gene (RNase H1SPBC.336.06c) cDNA was amplified by PCR from genomic DNA and ligated between the *Xho*I and *Bam*HI sites in pREP3X vector to obtain a thiamine repressible *rnh1* expression construct. Similarly, the construct *nmt1-rnh20*1 (a gift of Y. Murakami), encoding the subunit A of the three-subunit enzyme RNase H2 (SPAC4G9.02), was PCR amplified from genomic DNA and cloned between the restriction sites *Bam*HI and *Sal*I upstream of the *nmt1* promoter in the vector pREP1. All fission yeast media were prepared as described earlier (Moreno *et al,* 1991). The plasmids used in this study are listed in Supplementary Table5.

### Genetic Techniques

Sporulation was checked either microscopically or by staining the colonies with iodine vapours. Efficiently switching strains produce equal number of cells of opposite mating type, which mate to form zygotic asci that sporulate on minimal media. The spore cell wall contains a starchy compound that stains dark purple with iodine vapors. Thus, efficiently switching cells of an *h^90^* strain grow into colonies that stain dark purple with iodine while colonies of cells that switch inefficiently give light yellowish staining (Moreno *et al,* 1991). Determining the percentage of dark colonies provided a semi-quantitative measure of the switching efficiency. After iodine staining the colonies were photographed under Olympus Stereo Zoom Microscope.

Silencing status of any reporter gene was assessed by monitoring its expression using different plate assays. Expression of the *ura4* reporter was detected by dilution spotting, wherein 5μl of ten-fold serial dilutions of the overnight cultures of the required strains were spotted on non-selective plates, plates lacking uracil and those containing 5’FOA. FOA provided counter-selection to *ura^-^* cells. Thus, ura^+^ strains that grow well on ura minus but grow poorly on FOA plates, while ura-cells grow poorly on ura minus but grow well on FOA plates. Cell densities of all the cultures were normalized before serial dilution.

Expression of heterochromatic *ade6* was assessed by streaking the cells for single colonies on plates containing limited adenine. Cells expressing *ade6* produced light pink or white colonies on these plates and cells failing to express *ade6* gene produced dark red colonies. The fraction of colonies giving pink/white color provide a qualitative measure of the silencing defect. Similarly, the extent of silencing of the *his3* locus at subtelomeric locus, denoted as *his3-telo* (Cooper *et al*, 1997) was measured qualitatively by growth on plates lacking histidine.

### Cloning, site-directed mutagenesis, protein over-expression and purification

Lysine residues (KKK) at positions 242-244 in Swi6 were mutated to alanine (AAA) in the *nmt1-GFP-swi6^+^* expression construct (kindly gifted by Dr. Alison Pidoux) using the QuickChange site-directed mutagenesis kit (Stratagene), as per the manufacturer’s instructions, and confirmed by sequencing. The mutant is referred to as *swi6^3K→3A^*. The wild type full length Swi6 and Swi6*^3K→3A^* were tagged with GST at amino-terminus by cloning into pGEX-KG expression vector as BamHI-HindIII fragments. The vectors are listed in Supplementary Table S5.

### Expression and purification of GST-tagged Swi6 protein

The recombinant constructs containing GST, GST-Swi6, GST-Swi6^3K→3A^, GST-Swi6 -CD, GST-Swi6-CD-Hinge and GST-Swi6 (CSD) were transformed in to BL21(DE3) host cells. The transformed cells were cultured in the LB-Amp medium at 37°C. At mid-log phase, cells were induced with 1mM IPTG and incubated further at 25°C for 8hrs. For purification of GST-tagged proteins, 2-4 mg protein extract was added to 50 µl of Glutathione Sepharose beads (Cat. no.17527901, GE healthcare Ltd.) equilibrated with 1XPBS and incubated for 10-14 hours at 4°C. Purified proteins were collected after subjecting the mixture to elution for 2hrs at 4°C with 250 µl elution buffer containing 10mM reduced glutathione. Purification was confirmed by SDS-PAGE.

Expression of the *nmt1* promoter driven GFP tagged *swi6^+^* as well as *swi6^3K→3A^* gene was induced by removing thiamine from the medium of an overnight grown culture by washing the cells thoroughly and growing them further in medium lacking thiamine. It took 17-18 hrs of induction to achieve maximum expression level. Swi6 protein was detected by Western blotting with either anti-Swi6 (in-house, 1:10000), anti-GFP [Santa Cruz (sc-9996) 1:1000] or anti-GST [Santa Cruz (sc-138), 1:1000] antibodies. Wherever required, anti-α-tubulin antibody (Sigma, cat# T-9026; 1:4000) was used to probe the loading control. Alkaline phosphatase conjugated anti-rabbit (Sigma, cat# A-3562; 1:20,000) antibody was used, as per instructions. Western blots were developed using nitroblue tetrazolium (NBT) and bromo-chloro-indolyl phosphate (BCIP) as chromogenic substrates.

### Fluorescence Microscopy

10ml cultures of cells expressing the empty vector, *nmt1-GFP-swi6^+^* or *nmt1-GFP-swi6^3K→3A^* in *swi6Δ* background were harvested by centrifugation at 5000 rpm at RT. Cells were fixed in chilled 70% ethanol, rehydrated sequentially in 50%, 30%, 20% ethanol and finally in water. The pellets were resuspended in 50µl of 1X PBS. 5µl of this suspension was then visualized under fluorescence microscope. Similar method was followed for visualizing subcellular localization of chromosomally GFP-tagged Swi6 under the influence of over-expressed *rnh1* and *rnh201*.

### Generation of [γ-^32^P] labelled single stranded *in vitro* transcribed RNAs (ssRNA) and RNA-DNA hybrids

The protocol for synthesis and labeling of the single stranded RNA complementary to the heterochromatin-specific DNA sequences (Reihardt & Bartel., 2002; Djupedal *et al*, 2009) and the cognate RNA-DNA hybrids, is provided in Expanded View. Sequences of siRNAs A-L (Supplementary Table1) and I-XI (Supplementary Table S2) were as described (Reinhardt and Bartel, 2002; Djupedal et al, 2009). The template DNAs for synthesizing siRNAs in vitro are listed in Supplementary Table S3.

### Electrophoretic mobility shift assay (EMSA)

For EMSA, purified protein extract was mixed with ss siRNA or siRNA-DNA hybrid that was radiolabeled with [γ-^32^P] ATP with polynucleotide kinase in the binding buffer containing 20mM HEPES (pH 7.5), 50mM KCl, 2mM EDTA, 0.01% NP40, 1mM DTT, 5 units RNase inhibitor, 1 µg BSA and 5% glycerol along with ∼5 µg budding yeast total RNA or poly dI-dC as non-specific competitor, respectively, followed by incubation on ice. The samples were loaded on native TGE/ polyacrylamide (the tracking dyes Bromophenol blue and Xylene cyanol were not used in loading buffer as they interfere with the binding of proteins with nucleic acids). A negative control (without protein) was also loaded on the gel. 5X loading buffer containing both tracking dyes was added to negative control to track migration. For competition experiment, 0.1-10 pmol of ‘cold’ unlabeled RNAs and DNAs were used. The gel was dried in gel dryer and exposed to X-ray film or scanned in a phosphoimager.

The band intensities of the DNA-protein complex and free DNA for each lane were quantified using Scion Image software and the data were plotted according to a non-linear regression equation corresponding to one site specific-binding model as follows: Y = Bmax.X /(Kd+X), where Y represents the amount of protein-bound radioactivity, X, the protein concentration, Bmax the total change in radioactivity counts and Kd, the equilibrium binding constant (Heffler *et al*, 2012).

### Chromatin Immunoprecipitation (ChIP) Assay

Cells of appropriate strains were grown to an OD600 of 0.6 and crosslinked for 30 min with 3% formaldehyde at 18°C. Immunoprecipitation was performed with 1 µl anti-H3K9me_2_ (Upstate, 07-353) or 2 µl of polyclonal anti-Swi6 antibody per 400 µl reaction. Multiplex radioactive PCR amplifications of the immunoprecipitated chromatin samples, in the presence of [α-^32^P] ATP as the source of radio-labelled ATP in addition to non-radioactive dNTPs, were done using the following primers

ADE6F:5’-TGCGATGCACCTGACCAGGAAAGT-3’;

ADE6R: 5’-AGAGTTGGGTGTTGATTTCGCTGA-3’

URAF: 5’-GAGGGGATGAAAAATCCCAT-3’

URA4R: 5’-TTCGACAACAGGATTACGACC-3’

HIS3F: 5’-AGGTGTCCTTCTTGATGCCA-3’

HIS3R: 5’-CGAATTCCTGCTAGACCGAA-3’

dh For: 5’-GGAGTTGCGCAAACGAAGTT-3’

dh Rev: 5’-CTCACTCAAGTCCAATCGCA-3’

act1F: 5’-TCCTACGTTGGTGATGAAGC-3’

act1R: 5’-TCCGATAGTGATAACTTGAC-3’

The products are resolved on 4% 1X TBE polyacrylamide gel electrophoresis. The gel was exposed to the magnetic screen and scanned in Fuji Image Processor. The bands are quantified by densitometry from three independent sets of PCR using inbuilt MultiGauge Software. Data presented as mean SE. Data was analyzed through one-way ANOVA with Turkey’s post-hoc test where *** denotes p<0.001 when compared with vector and ns stands for non-significant.

### DNA-RNA hybrid Immunoprecipitation (DRIP) Analysis

Cells harvested from 500 ml log phase culture were washed with 1X PBS, resuspended in 2 ml PEMS (100mM PIPES, 1mM MgCl2, 1mM EDTA, 1.2 M sorbitol) supplemented with 20µl lyticase/zymolyase and incubated at 37°C for 30 min for spheroplasting. Spheroplasts were resuspended in 1.6ml TE containing 42µl of 20% SDS and 10µl of 10mg/ml Proteinase K and overnight at 37°C. Supernatant was extracted once with phenol:CIA and genomic DNA in the aqueous phase ethanol precipitated. 10µg of this DNA was digested overnight with Sau3AI at 37°C in 150 µl reaction. Following a phenol clean-up ethanol precipitated Sau3AI digested DNA was dissolved in TE without RNase A. One aliquot of the digested DNA was subjected to overnight RNase H (NEB cat#M0297) treatment at 37°C. Subsequently, the RNAse H treated samples were purified by phenol extraction and ethanol precipitation and dissolved in 50 µl TE mixed with binding buffer [10mM NaPO4 (pH 7.0), 0.14M NaCl, 0.05% TritonX-100]. 50 µl of both RNase H treated and untreated samples were saved as non-immunoprecipitated samples (NIPs). The DNA was immunoprecipitated overnight with 2 µl of anti-DNA-RNA hybrid mouse mono clonal antibody S9.6 (ATCC HB8730; ref 26) with protein G Sepharose beads and eluted in a buffer containing 50mM Tris (pH 8.0), 10mM EDTA, 0.5% SDS and 1.5 µg/ml Proteinase K at 55°C for 45 min. DNA from this complex was retrieved through phenol:CIA clean-up followed by ethanol precipitation. 2-3ng of this DNA was used for each RT-qPCR reaction set up with primers specific to the centromeric *dh* repeat region using SYBR Master Mix (Thermo Fisher) in QuantStudio3 (Thermo Fisher). Fold enrichment was calculated using ΔΔC_t_ method (Lival & Schmittgen, 2001).

## ACKNOWLEDGEMENTS

We thank Y. Murakami for the gift of the plasmids expressing rnh1 and rnh201 and R. Allshire for the strains with ade6 and ura4 reporters at the *cenI* locus.

