## Supplementary File for "An H3-K9-me-independent binding of Swi6/HP1 to siRNA-DNA hybrids initiates heterochromatin assembly at cognate dg-dh repeats in Fission Yeast"

### SUPPLEMENTARY SECTION

### Methods

#### a) *in vitro* transcription

An *in vitro* transcription protocol was adapted from Wilusz lab protocol (http://csu-cvmbs.colostate.edu/academics/mip/wilusz-lab/Pages/lab-protocols.aspx) with suitable modifications. The reaction was carried out by assembling nuclease free water, 1µl of DNA template (1µg/µl conc.), 4 µl of ribonucleotides mix (2.5mM each of ATP, GTP, CTP, and UTP) to a final concentration of 0.5mM, 2 µl of 10X RNA polymerase buffer and 1 µl of T7 RNA polymerase enzyme (Cat. no. AM2178, Ambion) were mixed and diluted to 20 µl with nuclease-free water and incubated at 37˚C for exactly 3 hours. The reaction was subjected to Phenol (pH 5.2)/Chloroform extraction, followed by ethanol precipitation.

#### b) Polyacrylamide gel elution of *in vitro* transcribed RNAs

To obtain ~500 ng of *in vitro* transcribed RNA, products of 10 reactions were pooled. The pellet obtained after precipitation was resuspended in 20 µl 1X formamide dye and loaded on 10- 20% polyacrylamide/8M urea gels depending on the size of the required transcript. Subsequently, the gel was stained with Syber gold nucleic acid stain (Cat. no. S11494, Invitrogen Life Technologies) for visualization under UV transilluminator. The RNA bands were located along the RNA marker (Cat. no. AM7778, Ambion), excised and frozen at -20$^{o}$C for 20 min. Thereafter, the gel pieces were crushed and resuspended in 1 ml of Crush and Soak buffer. The tubes were incubated at 37˚C and 700 rpm for 12-14 hrs on a thermo mixer. The supernatant was then subjected to ethanol precipitation.

#### c) Dephosphorylation of 5'ends of RNA

The ssRNA generated by *in vitro* transcription was dephosphorylated before labeling. The dephosphorylation reaction was set up with 500ng of ssRNA, Alkaline phosphatase calf intestinal enzyme, CIP (Cat. no. M0290, New England BioLabs Inc., USA), 1X CIP buffer, RNase inhibitor (Cat. no. AM2696, Ambion) and nuclease-free water up to 25 µl. The reaction was incubated at 37°C for 1 hour, phenol/chloroform extracted and ethanol precipitated.

#### d) 5'-end labeling of RNA

To perform 5'-end labeling of dephosphorylated RNA, a reaction mix containing 500ng of dephosphorylated RNA, T4 Polynucleotide kinase (PNK) enzyme (Cat. no. M0201S, New England BioLabs Inc., USA), 1X T4 PNK buffer, RNase inhibitor, [$\gamma$-$^{32}$P]-dATP (>3300 Ci/mmol) from Board of Radiation and Isotope Technology (BARC), Mumbai, and nuclease-free water up to 50μl was prepared and incubated at 37°C for 1hr. Unincorporated isotope was removed by passing through microspin G-50 columns (Cat. no. GE27-5330-01, GE Healthcare Ltd.). The eluent was stored at -70°C and used for performing RNA-EMSA.

#### e) RNA-DNA hybrid formation

For producing RNA-DNA hybrids, the ssRNA was annealed with equimolar amount of complementary DNA oligo in presence of 1X transcription buffer. The reactions were incubated at 85°C for 5 minutes to remove the secondary structures and then subjected to gradual cooling to 55°C. The reaction was incubated at 55°C overnight for annealing and checked on 10% native TBE gel before using for the EMSA experiments.

#### f) Synthesis of long RNAs

For *in vitro* synthesis of '*RevCen*' RNA, a long centromeric sequence shown in Djupedal *et al* (2009) was PCR amplified and cloned into the vector Litmus-28i. The clone containing the DNA sequence complementary to '*RevCen*' RNA was digested with *Hin*dIII to obtain a linear plasmid. The linearized plasmid was used as a template for *in vitro* transcription, as described earlier. The *in vitro* transcribed RNA was gel purified and subjected to alkaline phosphatase treatment followed by 5'end-labeling with [$\gamma$-$^{32}$P] dATP (10 μCi/μl, specific activity 3300 Ci/mmol) and used for EMSA experiments. The 'Cen100' transcript, as reported (Keller *et al*, 2002; SPAC15E1.04), was also synthesized and labeled by the same protocol. The radio-labelled oligonucleotides were purified by passing through a BioGel P-6 column (Biorad Inc.).

### g) RIP-Seq Analysis

Small RNA was isolated from immunoprecipitated samples from cells expressing GFP-Swi6$^{+}$, GFP-Swi6$^{3K\rightarrow3A}$ and TAP-Tas3. Cells from 500 ml cultures were harvested at log-phase (OD595= 0.5-0.7) were washed once with 10 ml ice-cold water and once with ice-cold STOP buffer (0.5% SDS, 5mM EDTA pH 8, 100 µg proteinase K) at 4°C. Cells were resuspended in (one-fifth the culture volume) HB buffer (25mM MOPS pH 7.2, 15mM MgCl2, 15mM EGTA, 1mM DTT, 1% Triton-X100, 10% glycerol) supplemented with PMSF (to a final concentration of 1mM) and protease inhibitor cocktail (PIC) (at 1:100 dilution) and dispensed as 1 ml aliquots into bead beater tubes. Ice-cold zirconium or acid-washed RNase-free glass beads were added upto 500µl mark, given 10-12 pulses of 1 min each alternated with 5 min incubations on ice. The lysate was centrifuged at 13500 rpm at 4°C for 1 hr. The clarified supernatants were removed to fresh tubes and mixed with 100% glycerol to a final concentration of 10%. The expression of the protein of interest can be checked at this point by Western analysis (as described previously). The extracts were kept at -80°C until further use. For immunoprecipitating TAP tagged proteins, commercially available TAP antibody conjugated agarose beads (Santa Cruz, sc-32319 AC) were used and no coupling was needed. These beads were directly used after equilibration with IP buffer. 2µl of anti-Swi6 antibody was conjugated to 50 µl protein-A Sepharose beads (Sigma, P9424-1ml) in a 500 µl 1X PBS containing coupling reaction for 16hrs at 4°C with continuous mixing. The antibody bound beads are then washed twice with 500µl of 0.2M borate buffer (pH 9) and subsequently incubated in 500µl of 0.2M respective borate buffer containing dimethylpimelimidate (DMP) at a concentration of 5mg/ml for 30 minutes at RT with mixing. DMP-mediated coupling was stopped by washing the beads once with 500µl of 0.2M ethanolamine (pH 8.0). The residual DMP was quenched by incubating the beads in 500µl 0.2M of ethanolamine (pH 8.0) for 2 hrs at RT. The beads were then washed once with 1X IP buffer [50mM Tris-Cl (pH 7.5), 150mM NaCl, 0.5% NP40 and 0.5mg/ml BSA). Approximately 3 mg of crude extract was bound in the presence of 1X IP buffer for 16hrs at 4°C with mixing. Beads were then washed once with IP wash buffer [50mM Tris-Cl (pH 7.5), 150mM NaCl, 0.1% NP40 (or 0.1% Tween-20) and 1mM EDTA (pH 8.0)]. To each 50 µl beads loaded with immunoprecipitate, 200µl each of LETS buffer [100mM LiCl, 10mM

EDTA (pH8.0), 10mM Tris (pH 7.4), 0.2% SDS] and phenol (pH 5.2) were added. Samples were vortexed gently with intermittent ice incubation (30 sec on, 30sec off) followed by 1 hr at 37°C. The beads are then centrifuged at RT for 5min at 12000rpm. The collected supernatants were extracted once with equal volumes (~400µl) of phenol (pH 5.2): chloroform (1:1). RNA in the upper aqueous phase was precipitated with 20µg glycogen, one-tenth volume of 5M LiCl (40µl) and 2.5 volume of chilled absolute ethanol (1100µl) at -80ºC for 16hrs. Pellet was washed with 300µl chilled 80% ethanol and dissolved in 250µl of RNase-free water. To the dissolved RNA, 250µl PEG precipitation solution (20% PEG (MW8000) in 2M NaCl) was added. Contents were incubated on ice for 30 min and centrifuged at 12000rpm for 20 min at 4ºC. The supernatants were transferred to fresh tubes while the pellets were first dissolved in TE (pH 8.0), ethanol precipitated, washed with 80% ethanol, dried and resuspended in RNase-free water. This contains the population of long RNA while the supernatant isolated after PEG precipitation contains the small RNAs. Supernatant obtained after PEG precipitation was extracted once with phenol (pH 5.2): chloroform and small RNA present in the aqueous layer was precipitated using LiCl-ethanol system as earlier. The isolated small RNA was checked by running on a 15% 7M denaturing urea gel. This RNA was subjected to Northern blotting with probes complementary to *dh* sequences of the *S. pombe* centromeric repeat regions. Alternatively, they were used as riboprobes for Southern blots of PCR products generated from the centromeres and the mating type loci of fission yeast, like, *dh, dhk* and *dg*; *act1* was taken as a negative control. Finally, one set of the isolate RNA was subjected to next generation sequencing.

The NEBNext® Multiplex Small RNA Library Prep Set for Illumina based system was used for 3' adaptor ligation, cDNA synthesis, 5' adaptor ligation and PCR, followed by size selection and sequencing using the Illumina platform was performed by Bionivid Technology Pvt. Ltd., Bangalore.

The sequencing data was analyzed using bowtie2 and R-packages as detailed in the following section.

### h) RIP-seq data processing

Gene annotations and reference genome of fission yeast were downloaded from https://www.pombase.org/ . Reference genome was indexed using 'bowtie2-build' * http://bowtie-bio.sourceforge.net/bowtie2/index.shtml). We trimmed the 5' and 3' Illumina small RNA-seq adapters sequences (GUUCAGAGUUCUACAGUCCGACGAUC and

TGGAATTCTCGGGTGCCAAGG, respectively) using 'cutadapt' (https://cutadapt.readthedocs.io/). Duplicates were removed using 'rmdup' utility of SeqKit package (https://bioinf.shenwei.me/seqkit). The clean reads were mapped onto the indexed reference genome using bowtie2 with '–sensitive' setting. The quality of the reads was assessed using FastQC (https://www.bioinformatics.babraham.ac.uk/projects/fastqc/) at each step. We binned the raw reads at 5kb (for boxplot) and 200 bp (for line plot) resolutions and quantile normalized the read counts across samples using the function 'normalize.quantiles' of 'preprocessCore' R-package (https://www.bioconductor.org/packages//2.11/bioc/html/preprocessCore.html). The centromere annotations were obtained from https://www.pombase.org/status/centromeres. The telomeric and sub-telomeric regions were defined as 0-50Kb and 50-100Kb regions from the chromosome ends. The read-counts for *swi6*$^{+}$ and *swi6*$^{3K\rightarrow 3A}$ samples were divided by those of control ('vector') sample. We calculated the log2 ratio of read counts of *swi6*$^{3K\rightarrow 3A}$ to *swi6*$^{+}$ for plotting purpose. Sequence and annotations for mating-type region were obtained from https://www.ncbi.nlm.nih.gov/nuccore/FP565355.1. The reads were independently mapped onto mating-type sequence following the same procedure as earlier and processed at 200 bp resolution. We further identified the 5kb bins that had at-least one of the sequence motifs known to interact with Swi6 (Table X) using in-house script. P-values were calculated using two-tailed Mann-Whitney U tests on R.

The quality scores of RIP-Seq data are shown in Supplementary Table S7.

### i) ChIP-seq data processing

We downloaded the 'sra' files of the ChIP-seq experiments, such as H3K4me3, H3K36me3, H3K9me2, H3K9me3, RNA PolII and Swi6 from GSE83495. We converted the files to ‘fastq’ format using ‘fastq-dump’ of NCBI SRAtoolkit. The reads were then mapped on the indexed genome reference using bowtie2 with default parameters. We converted the ‘sam’ files into ‘bam’ files using ‘samtools’ and ‘bam’ to ‘bed’ using ‘bedtools’ with default parameters. We binned the raw reads at 5kb resolution and labeled bins appropriately when their coordinates mapped into telomeres, centromeres and mating region. We plotted the density of read counts for the bins with RIP-seq log2 ratio (mutant to wild-type) > +1 or < -1. The p-values were calculated using Mann Whitney U tests.

#### j) Gene expression analysis

FPKM values for WT pombe strain were downloaded from GSE104546 and the average gene expression was calculated for each 5kb bin. We plotted the distribution of average gene expression values for the bins with RIP-seq log2 ratio (mutant to wild-type) $\geq$ +1 or $\leq$ -1. The p-value was calculated using Mann Whitney U test.

**Supplementary Figure S1**. **Mapping of the siRNA sequences to the *dg-dh* regions.**

A In *cenI*,

B In *cenH* in the mating type locus and

C In the *tlh1-tlh2* genes.

**Supplementary Figure S2**. Pairwise alignment of different siRNAs used in this study.

**Supplementary Figure S3**. **Experimental design for synthesis of siRNAs in vitro.**

A The double-stranded template, wherein the top DNA strand comprised of the T7 promoter sequence fused to DNA sequence cognate to the siRNA sequences, is transcribed in presence of ribonucleotides and T7 RNA polymerase to synthesize the respective siRNA. Red bar denotes the T7 promoter and green bar the DNA sequence cognate to the siRNA sequence.

B The full length and domains of Swi6: CD, CD-hinge and CSD, expressed as GST fusion.

C The purified proteins shown in B visualized by SDS-PAGE.

**Supplementary Figure S4. Binding of Swi6/HP1 to the 'E-For' siRNA and competition for binding by sequences I-IX (Djupedal et al, 2009).**

A EMSA assay was performed for binding of Swi6 to the radiolabelled 'E-For' RNA in presence of excess of unlabeled 'cold' RNAs with sequences of E and I-IX.

B Binding to 'E-For' RNA could also be competed out by excess of 'V' RNA that maps to the subtelomeric genes, *tlh1* and *tlh2* (Figure 1).

**Supplementary Figure S5. Consensus binding sequence for Swi6.**

The sequences, labeled B, D, E and K were aligned according to the software developed by Bailey et al (2009)..

**Supplementary Figure S6. Determination of affinity constant for binding of Swi6 at multiple binding sites in the D-For RNA.**

A. EMSA was performed to detect binding of GST and increasing concentration of GST-Swi6 with radiolabeled D-For RNA. The reactions were resolved by electrophoresis and subjected to autoradiography.

B Bands shown in A were quantitated by densitometry and plotted.

**Supplementary Figure S7. The siRNA precursor RNA 'RevCen' and 'Cen100', fail to show binding to Swi6 in EMSA assay.**

A. The EMSA assay was performed with the radiolabeled 'RevCen' RNA
B. The EMSA assay was performed with the radiolabeled 'Cen100' RNA.
   EMSA assay was performed by incubating the radiolabeled RNA with GST or GST-Swi6, and subjecting to electrophoresis in 8% acrylamide gel, as in Figure 1.

**Supplementary Figure S8. Specificity of binding of Swi6 to the *dg.dh* repeat sequences *in vivo*.**

A Expression of GFP-Swi6 and GFP-Swi6$^{3K\rightarrow 3A}$ in *swi6Δ* cells. Extracts prepared from cells of *swi6Δ* strain expressing the control vector, GFP-Swi6 and GFP-Swi6$^{3K\rightarrow 3A}$ were subjected to western blotting with anti-GFP (upper panel) and $\alpha$-tubulin (lower panel) antibody.

B Slot blot hybridization. RNAs extracted from cells expressing TAP-tagged Tas3 immunprecipitated with anti-CBP antibody and those expressing GFP-tagged Swi6 or GFP-tagged Swi6$^{3K\rightarrow 3A}$, immunoprecipitated with anti-GFP antibody, were radiolabeled at the 5'-end with [$\gamma^{32}$P] ATP and T4 polynucleotide kinase and used as a probe for slot blots on which equal amounts of the DNAs (100ng) corresponding to *act1*, *dg*, *dh* and *dhk* (*cenH*) were blotted. The bound signal was visualized by autoradiography.

C The size-fractionated radiolabeled siRNA samples from indicated strains were subjected to electrophoresis in 8% acrylamide/7M urea gels and visualized by autoradiography. In parallel, unlabeled samples were similarly processed blotted and probed with radiolabeled snoRNAs (lower panel), followed by autoradiography.

D No bidirectional transcription is observed in cells expressing *swi6*$^{3K\rightarrow 3A}$ mutant. RNAs extracted from wt, *dcr1Δ*, and *swi6Δ* strain harboring the empty vector, or vector expressing GFP-Swi6 or GFP-Swi6$^{3K\rightarrow 3A}$ were subjected to RTPCR for the Forward strand and Reverse strand for the *dh* and *act1*. The PCR products were visualized by agarose gel electrophoresis and staining with ethidium bromide.

**Supplementary Figure S9.**

A Chromosomal tracks of RIP-seq fold change (log2 *swi6*$^{3K\rightarrow 3A}$/*swi6*$^{+}$ ratio) at and around centromeres of chr-II and chr-III at 5kb (top) and 200bp (bottom) resolutions.

B RIP-seq fold change fold change at right telomere of chr-II (200b resolution). Locus annotations are given at the bottom of the plot.

**Supplementary Figure S10. The mutant *swi6*$^{3K\rightarrow 3A}$ is defective in heterochromatin localization.**

A Transformants of *swi6Δ* strain harboring empty vector, vector expressing GFP-Swi6 and GFP-Swi6$^{3K\rightarrow 3A}$, were visualized by confocal microscopy. Arrowheads indicate the spots (normally three in over 65% of cells) where GFP Swi6 is localized (middle panel, right) but not in case of GFP-Swi6$^{3K\rightarrow 3A}$.

B Quantitation of per cent cells containing one, two or three fluorescent spots of GFPSwi6, while in case of GFPSwi6$^{3K\rightarrow 3A}$, fewer cells (~20%) contain 3 spots, and more cells have two or one spot or diffuse fluorescence.

C	Schematic representation of the mating-type locus, indicating the insertion of *ura4* reporter at centromere distal location with respect to the *mat2P*. Also show are the homology boxes H2 and H1 at all three loci: *mat1*, *mat2* and *mat3* and H3 box at *mat2* and *mat3* loci. *REII* represents the repression element II and Δ sign indicates deletion of the *REII* site.

D	Spotting assay as a qualitative measure of the level of expression of the *ura4* reporter. Cells of *swi6Δ* strain with the genotype shown in (C) transformed with empty vector, GFP-Swi6 and GFP-Swi6$^{3K\rightarrow 3A}$, were serially diluted and spotted on plates lacking leucine (for plasmid selection) and those lacking leucine and containing FOA. The cells were allowed to grow for 4 days at 30$^{o}$C and photographed.

E	Swi6$^{3K\rightarrow 3A}$ fails to complement the switching defect in the $h^{90}$ *swi6*Δ mutant. Cells of indicated transformants of *swi6*Δ strain were spotted on minimal medium lacking iodine. After 4 days' growth at 30$^{o}$C, the colonies were stained with iodine and photographed.

**Supplementary Figure S11. Expression of *rnh1* abrogates the heterochromatin localization of GFP-Swi6**.

Cells expressing chromosomally tagged GFP-Swi6 were transformed with empty vector, and vector expressing *rnh1* and *rnh201* under the control of *nmt1* promoter, which is repressed in presence of thiamine. Cultures of the transformants grown in presence (+; repressed) and absence (-, derepressed) of thiamine were visualized by confocal microscopy. Representative data shown in Figure 4C.

**Supplementary Figure S12. RNA-DNA hybrid formation is essential for heterochromatin silencing and Swi6 localization.**

A	Schematic representation of mating type locus, showing the deletion of the *mat3*-linked repression element *REIII* (denoted by the Δ sign) and insertion of the *ade6* reporter distal to *mat3*.

B The silencing of the *ade6* reporter is abrogated by overexpression of *rnh1* but not *rnh201*. Transformants of the indicated strain were streaked on selective plates containing limiting amount of adenine. The plates were grown for 4 days at 30$^o$C and photographed.

C The fraction of transformant colonies giving pink/white colour were counted and plotted.

D ChIP assay was performed to quantitate the enrichment of Swi6 at the *mat3::ade6* locus in the transformants shown in (B).

E Schematic representation of the mating type locus *mat2*, indicating the deletion of repression element *REII* (denoted by the sign Δ) and insertion of *ura4* reporter at centromere distal location.

F Overexpression of *rnh1* but not *rnh201* causes loss of silencing at the *mat2* locus as well, as indicated by growth of colonies that give dark staining with iodine (Materials and Methods).

G Fraction of transformants with the different vectors giving dark staining with iodine was plotted.

H Delocalization of Swi6 at *mat2* locus upon transformation with *rnh1*. Level of Swi6 enrichment at the *mat2*-linked *ura4* locus in different transformants was plotted.

**Supplementary Figure S13. Enrichment densities**.

A Swi6

B RNA PolII

C H3K36me3

D Gene expression levels (FPKM)

Enrichment densities of A-D shown in the 5Kb genomic bins exhibiting 2-fold increase (log*swi6*$^{3K\rightarrow 3A}$/*swi6*$^{+}$ ≥1, orange) or 2-fold decrease (log2 *swi6*$^{3K\rightarrow 3A}$/$^{swi6+}$ ≤-1, blue) in siRNA Enrichment in *swi6*$^{3K\rightarrow 3A}$ mutant when compared with that of WT strain. P-values were calculated using Mann-Whitney U tests.

**Supplementary Figure S14. Log2 ratio (*swi6*$^{3K\rightarrow 3A}$/*swi6*$^{+}$) of RIP-seq profile at centromere of *chrI* and the flanking regions at 200 bp resolution.**

The thymidylate kinase locus (SPAC15E1.04) is highlighted in yellow colour. The zoomed image shows normalized RIP-seq signals at thymidylate kinase in *swi6*$^{+}$ and *swi6*$^{3K\rightarrow 3A}$ strains.

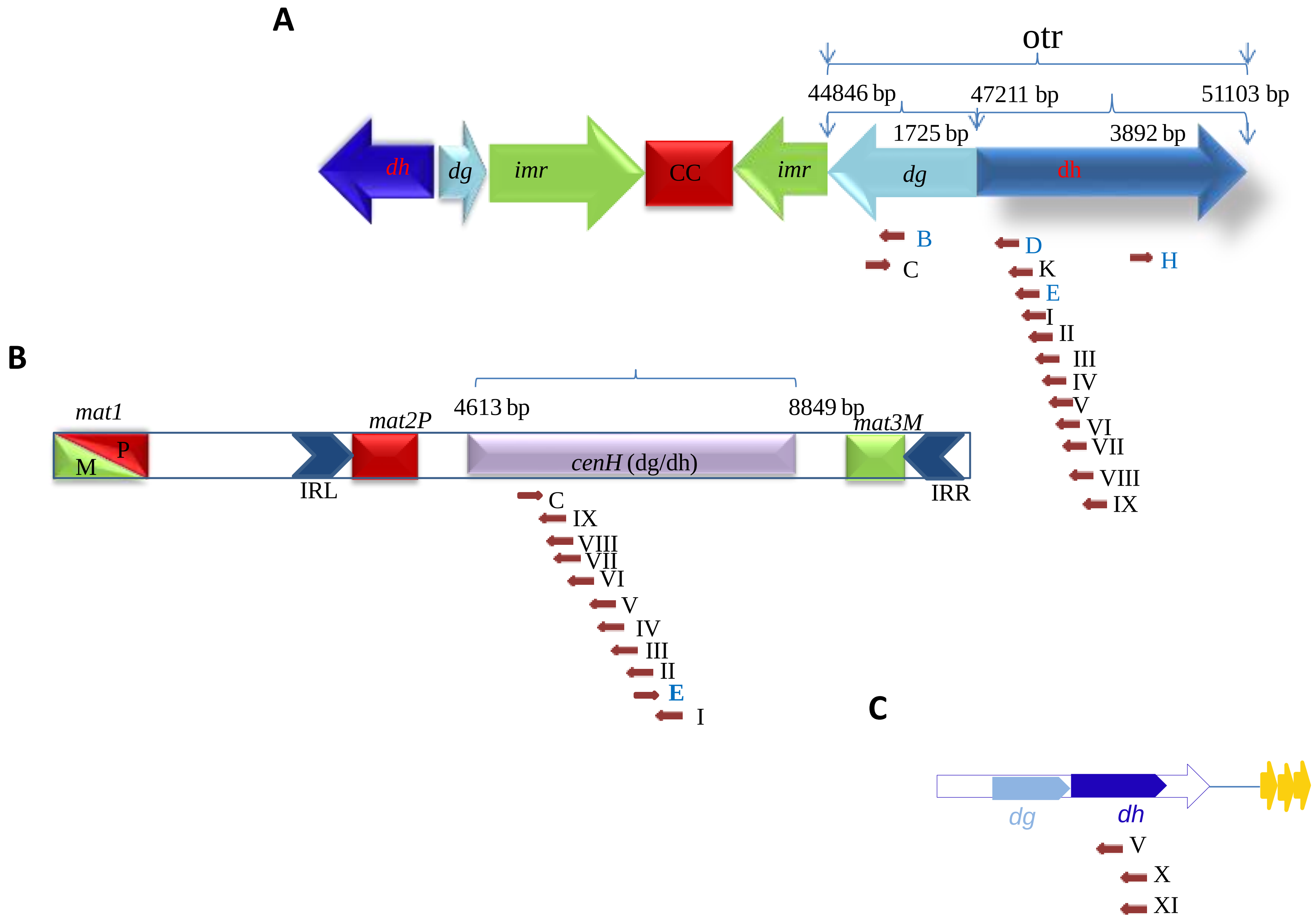


Supplementary Figure S1

```
Group 1      D    --UGGAUUAAGGAGAAGCGGUA--
'D' & 'J'    J    UCUACCUCAGCAGUCCUUGGAAA
                    *   *  *         ** *

Group 2      F     UGCGCAACUCCUGCUUAUCGUC
'F' & 'H'    H     UGAGCAUAUCCUAAUGACAGUA
                   ** ***  ****  * *  **

Group 3      I    ----UGCCUAUUUAUACAUUUCCC-
'I' & 'K'    K    UGUGUCCAUAUCCAUGCUGUGUCCA
                      * * ***  ** *  *  **

Group 4      E     ---ACAAGUGAUAAGAGUAGGUG
'E', 'G'     G     --UACAAGAUAUAGCGCCACACU
& 'L'        L     UAAACAACUUGCAAUAUCUGCCA
                      ****         *
```

RNA sequences from 'dh' repeat region :**D, E, H** and **K**
RNA sequence from 'dg' repeat region: **B**

Supplementary Figure S2

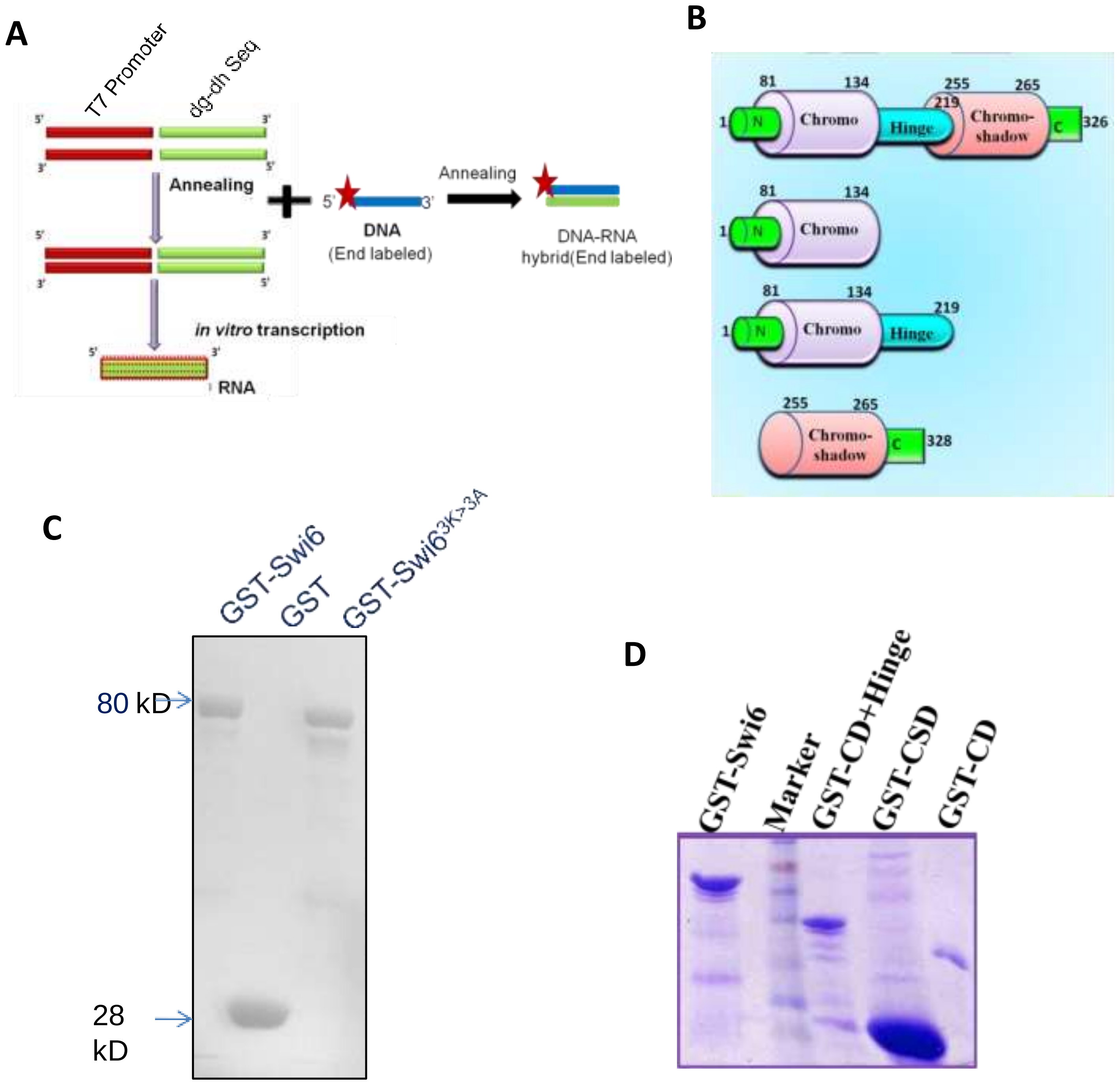


Supplementary Figure S3

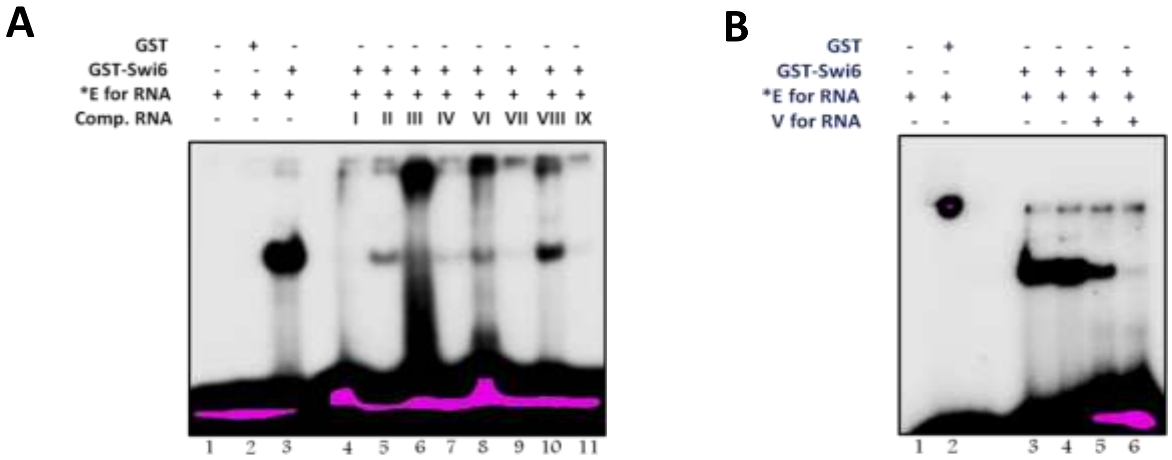


Supplementary Figure S4

### Discovered Motifs

**1.**

***E*-value:** 1.9e+002 **Site Count:** 4 **Width:** 10

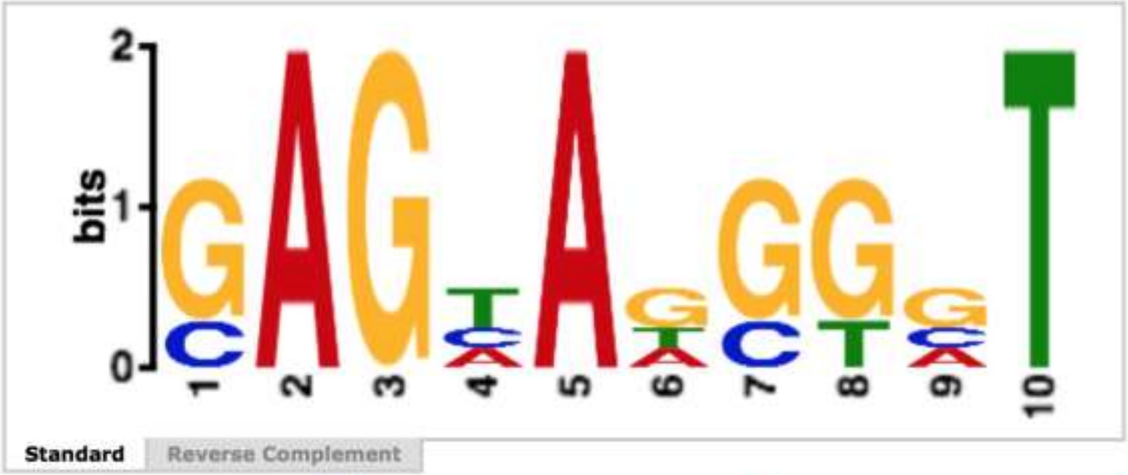


**Log Likelihood Ratio:** 37 **Information Content:** 13.1 **Relative Entropy:** 13.2 **Bayes Threshold:** 3.70044

| Name | Strand | Start | *p*-value | Sites |
|---|---|---|---|---|
| 3. E | + | 12 | 8.38e-6 | CAAGTGATAA GAGTAGGTGT |
| 1. B | + | 7 | 1.69e-5 | AATGCG GAGTAAGGCT AATCACGGTA |
| 2. D | + | 10 | 2.06e-5 | TGGATTAAG GAGAAGCGGT A |
| 4. K | - | 10 | 1.05e-4 | TGGACA CAGCATGGAT ATGGACACA |

Supplementary Figure S5

A

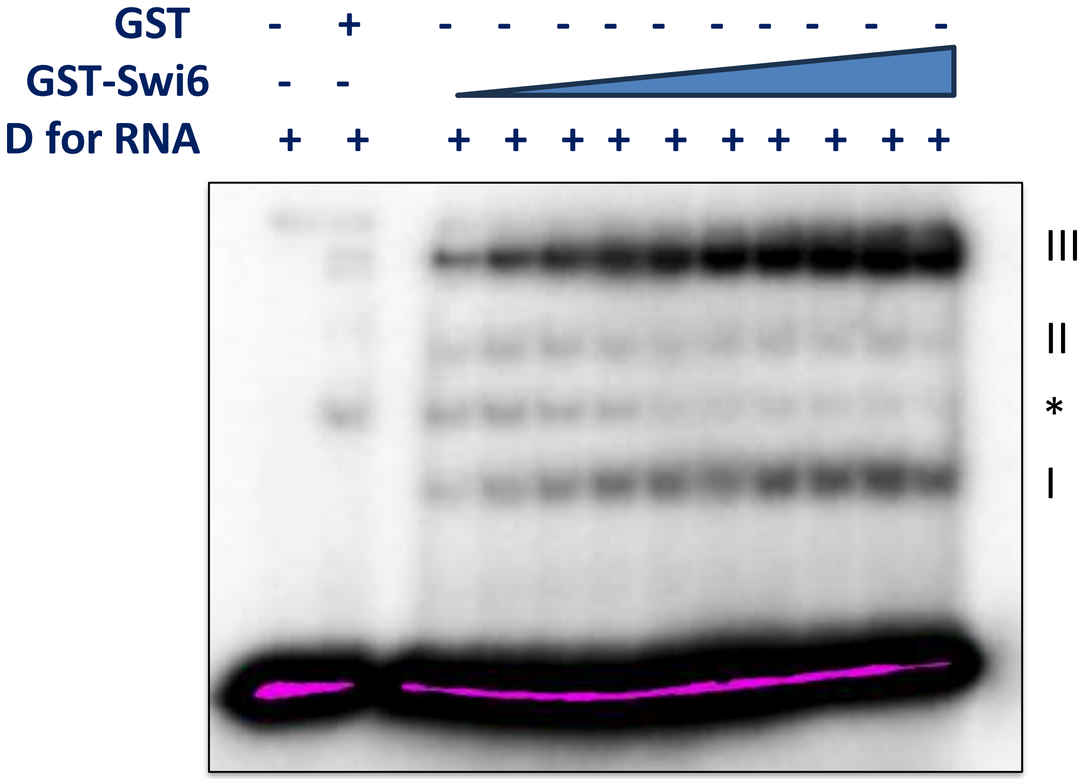
GST
GST-Swi6
D for RNA
III
II
*
I

B

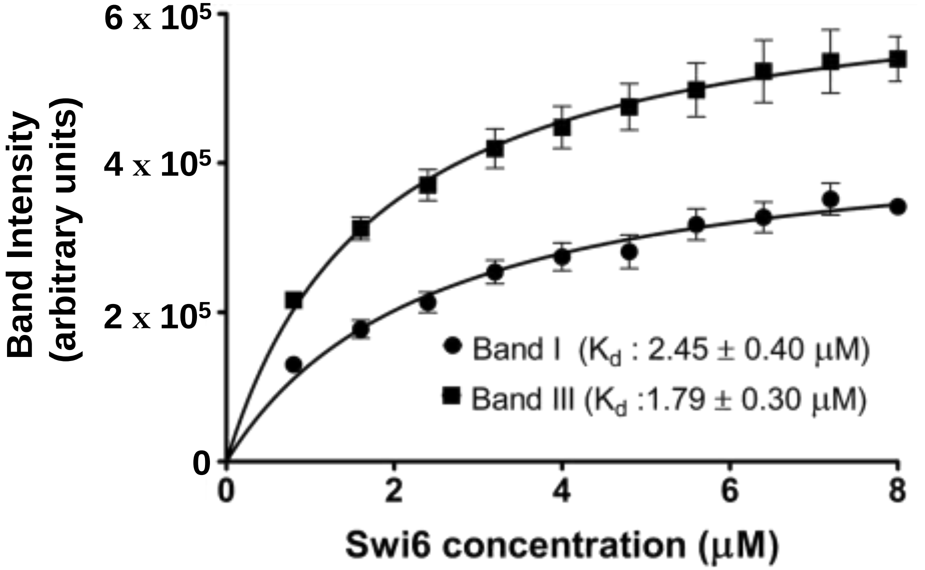
6 x 10⁵
4 x 10⁵
2 x 10⁵
0
Band Intensity (arbitrary units)
0
2
4
6
8
Swi6 concentration (μM)
Band I (Kd : 2.45 ± 0.40 μM)
Band III (Kd :1.79 ± 0.30 μM)

A

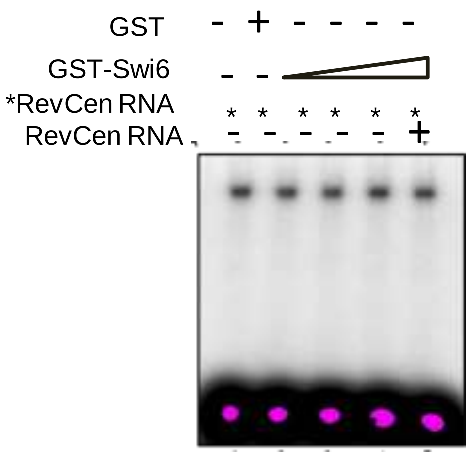


B

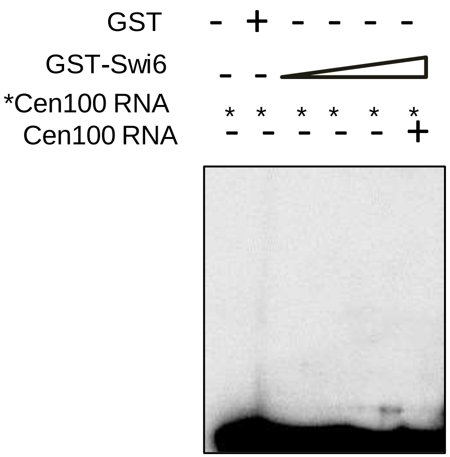


Supplementary Figure S7

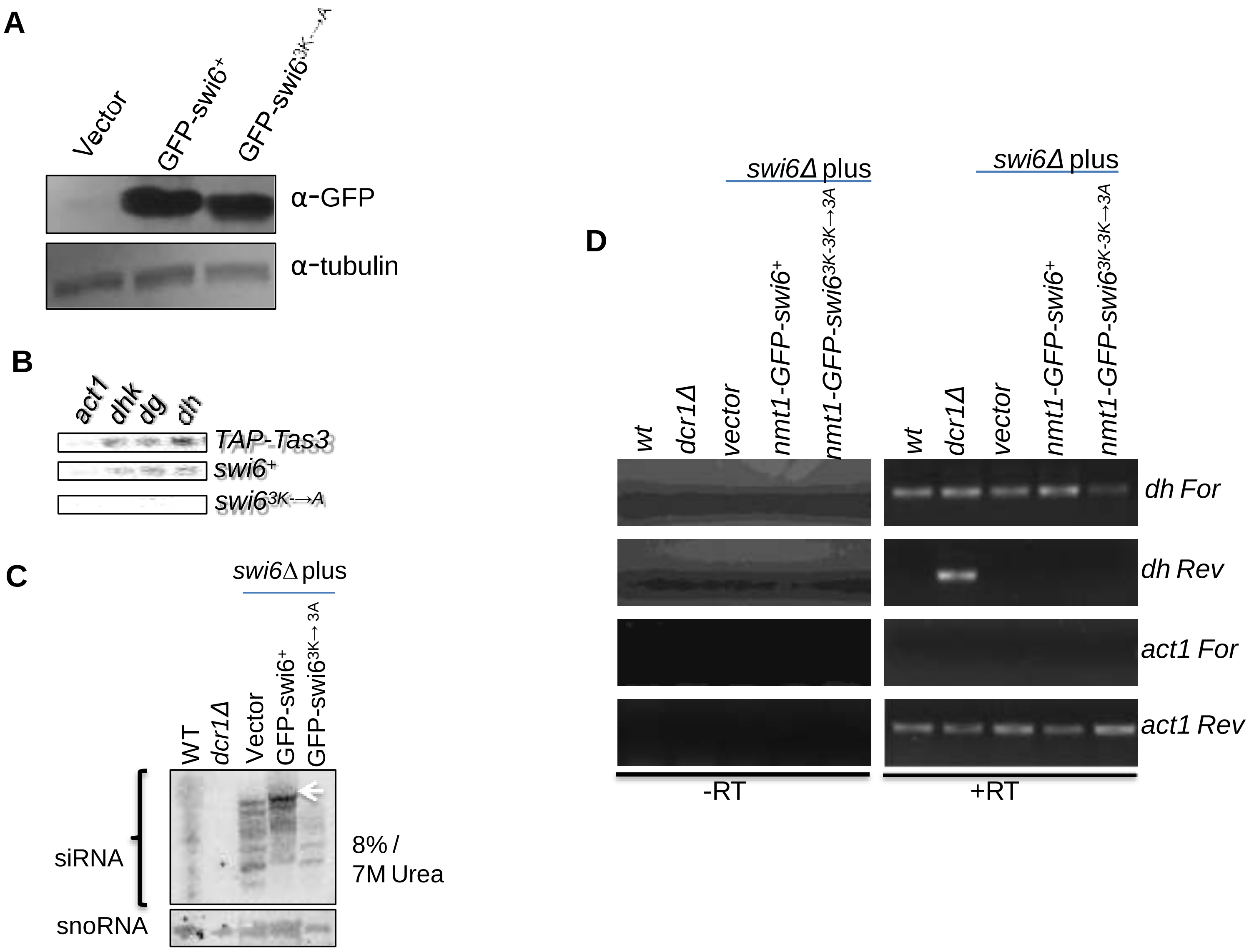


Supplementary Figure S8

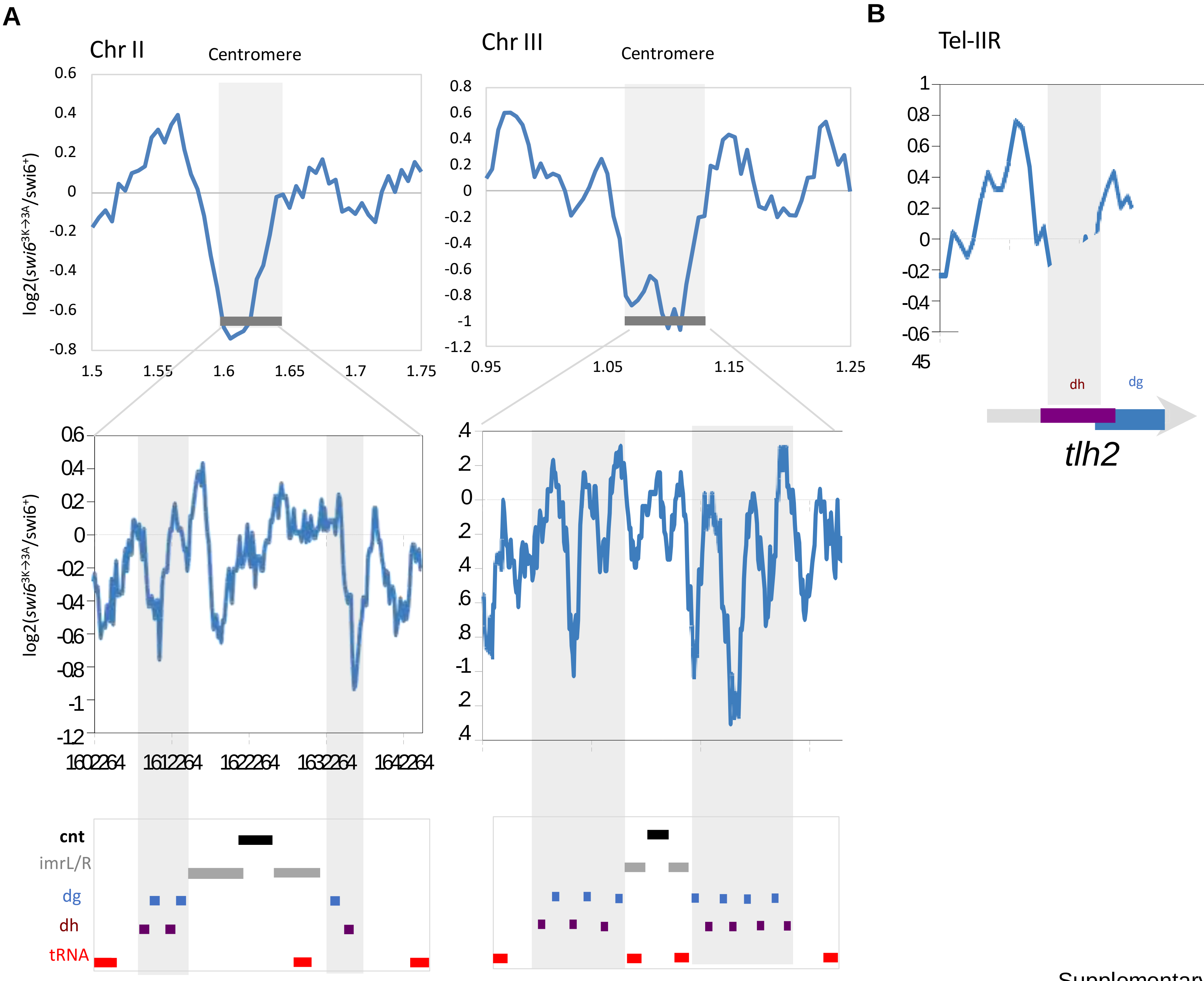


Supplementary Figure S9

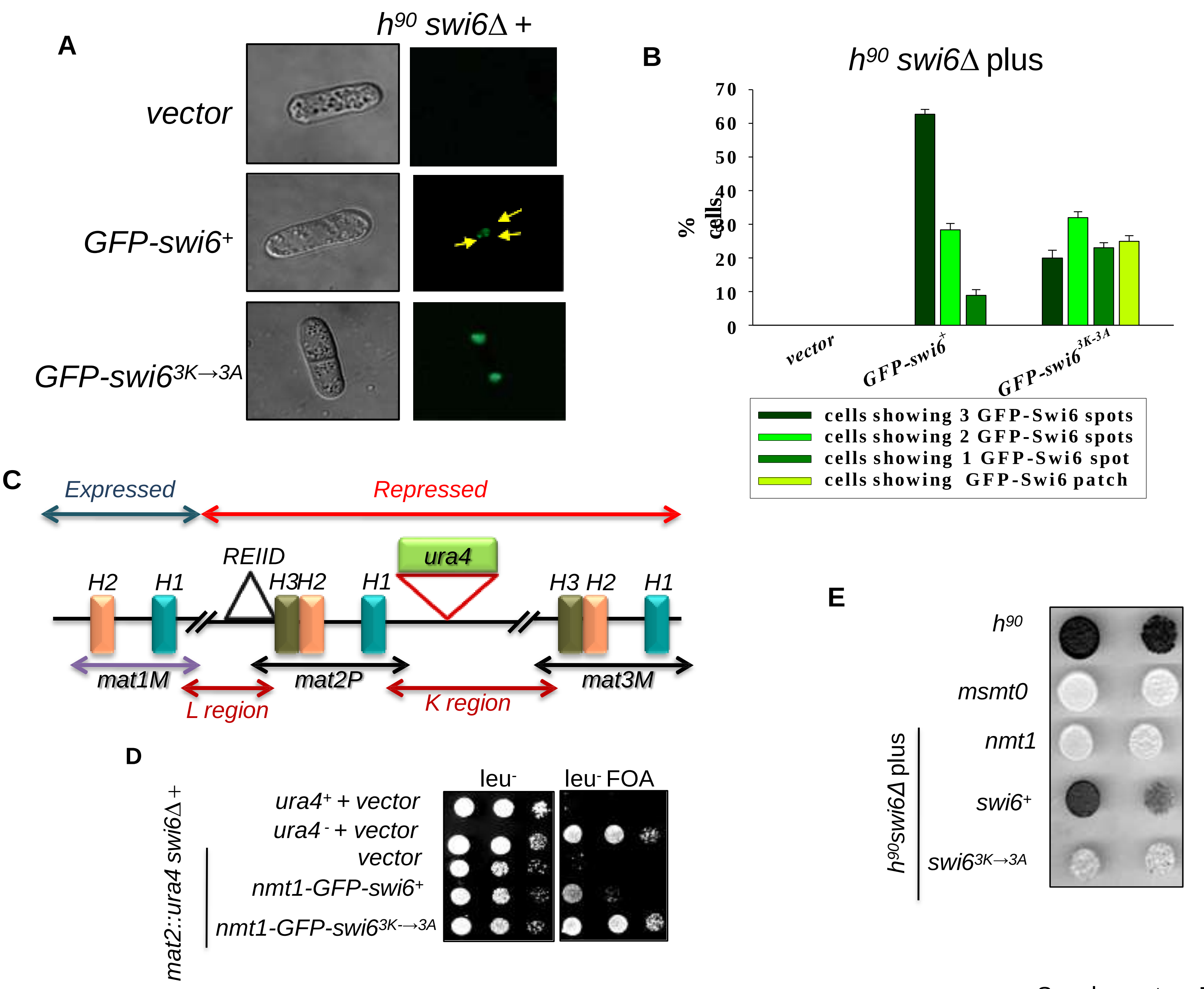


Supplementary Figure S10

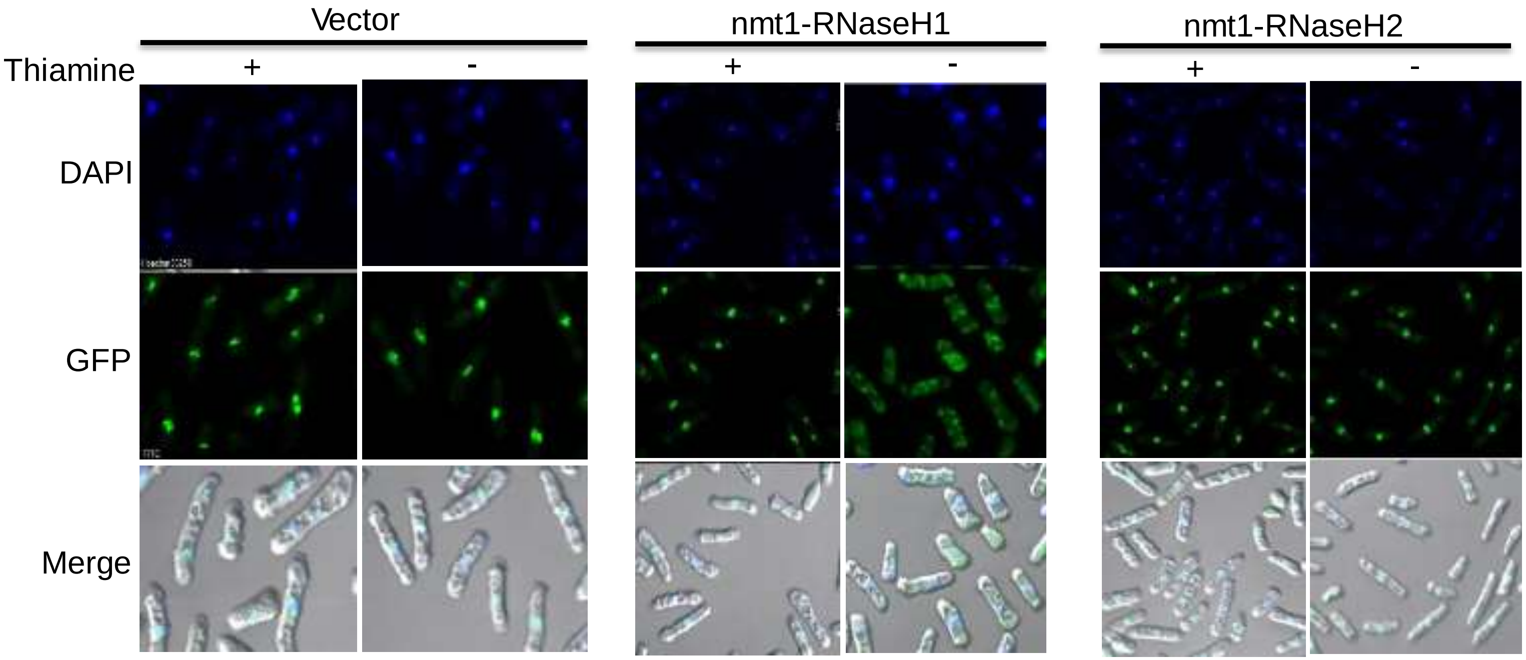


Supplementary Figure S11

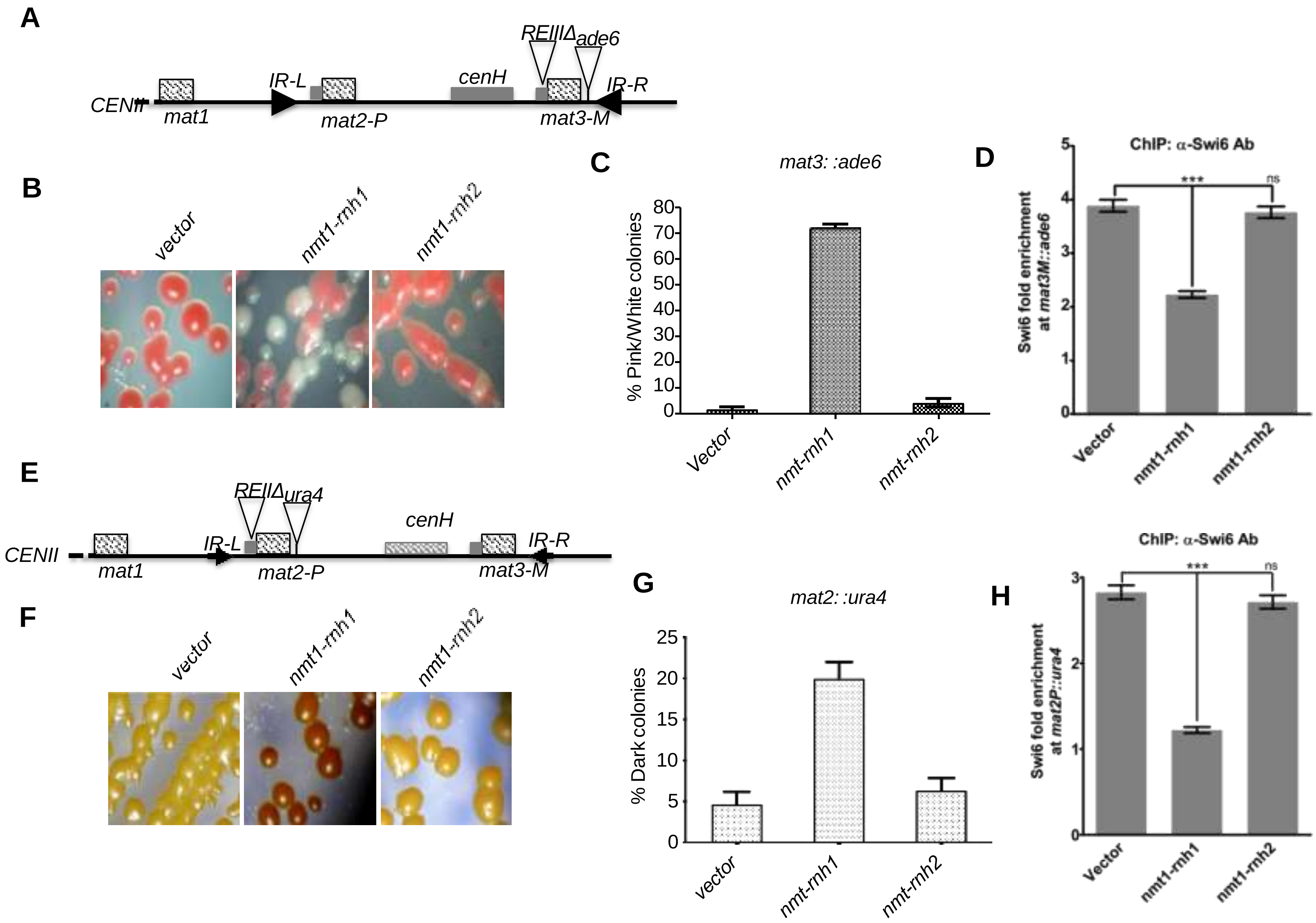


Supplementary Figure S12

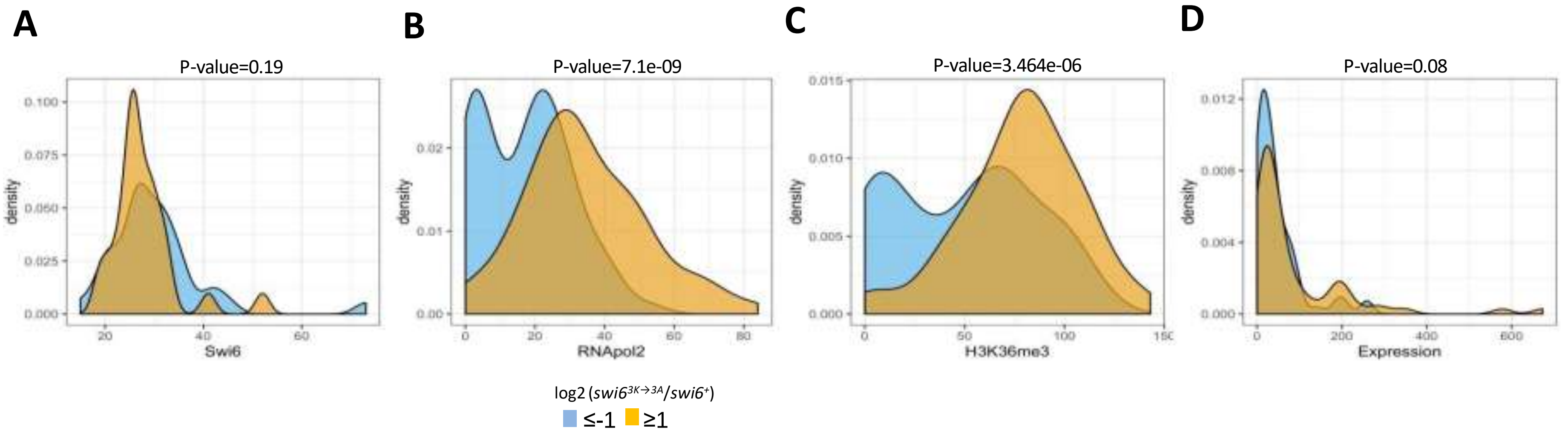


Supplementary Figure S13

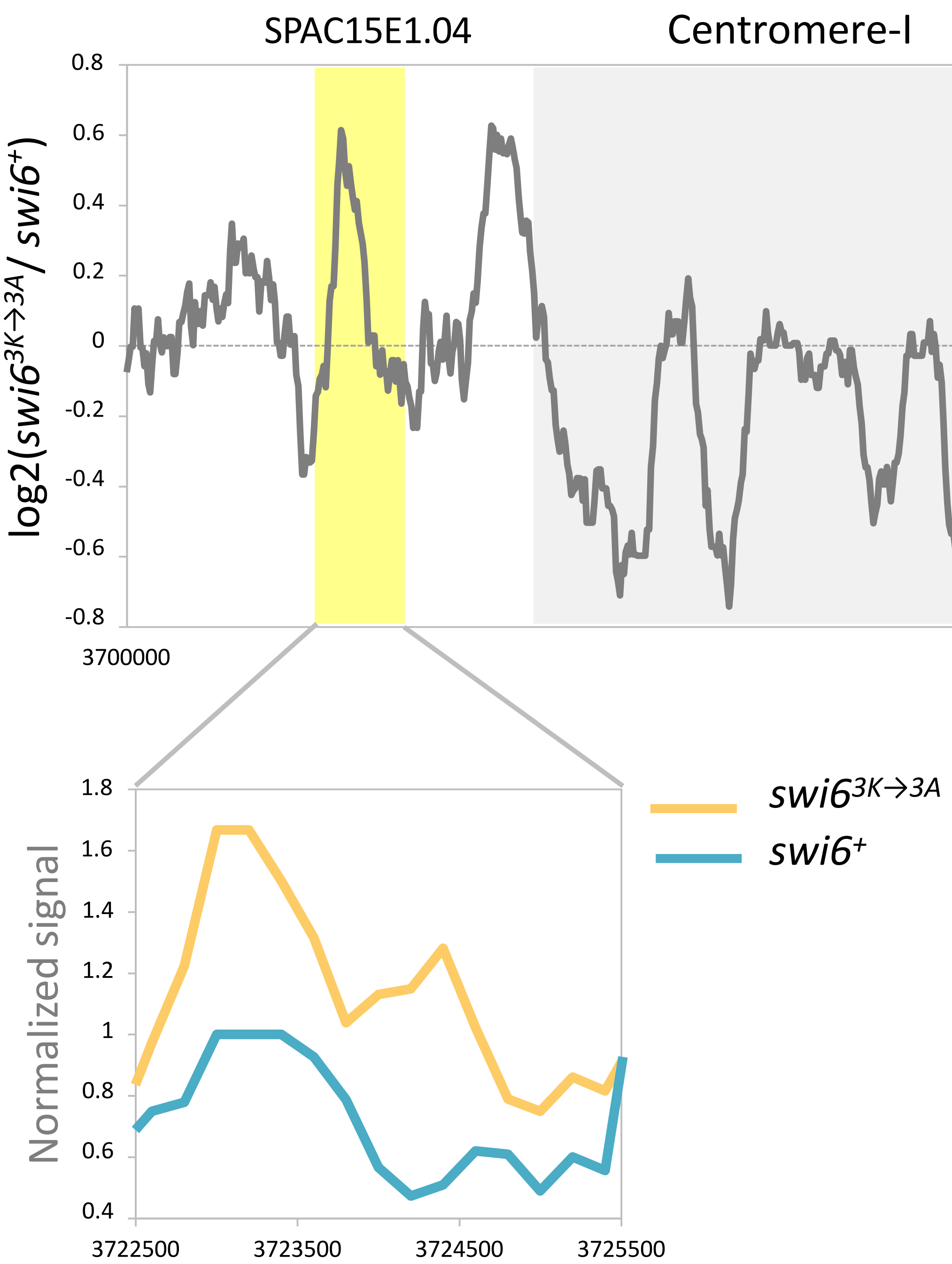


Supplementary Figure S14
